# MiR-7a-Klf4 axis as a regulator and therapeutic target of neuroinflammation and ferroptosis in Alzheimer’s disease

**DOI:** 10.1101/2025.03.24.644978

**Authors:** Madhu Ramesh, Thimmaiah Govindaraju

## Abstract

Neuroinflammation and ferroptosis significantly contribute to neuronal death in Alzheimer’s disease (AD) and other neurodegenerative disorders. MicroRNAs (miRNAs) are crucial regulators of these pathological processes. We employed transcriptomic analysis in an APP/PSEN1 Tg AD mouse model to identify dysregulated miRNAs and construct a miRNA-mRNA-pathway network. We discovered increased miR7a expression in the AD brain, targeting Krüppel-like factor 4 (Klf4), a transcriptional factor implicated in Aβ oligomer-induced neuroinflammation and RSL3-induced neuronal ferroptosis. Elevated Klf4 levels in AD mice brains suggest its involvement in AD pathology. The miR-7a mediated silencing of Klf4 alleviates neuroinflammation by modulating NF-κB, iNOS, and NLRP3 pathways, and inhibition of ferroptosis by targeting labile iron levels, GPX4, Nrf2 pathway, and mitochondrial damage. These findings highlight the neuroprotective role of miR-7a and its potential as RNA therapeutic. Pharmacological targeting of the miR-7a-Klf4 axis with blood-brain-barrier (BBB)-permeable compound effectively mitigates neuroinflammation and ferroptosis, suggesting the miR-7a-Klf4 axis as a novel therapeutic target for AD.

**GRAPHICAL ABSTRACT:** 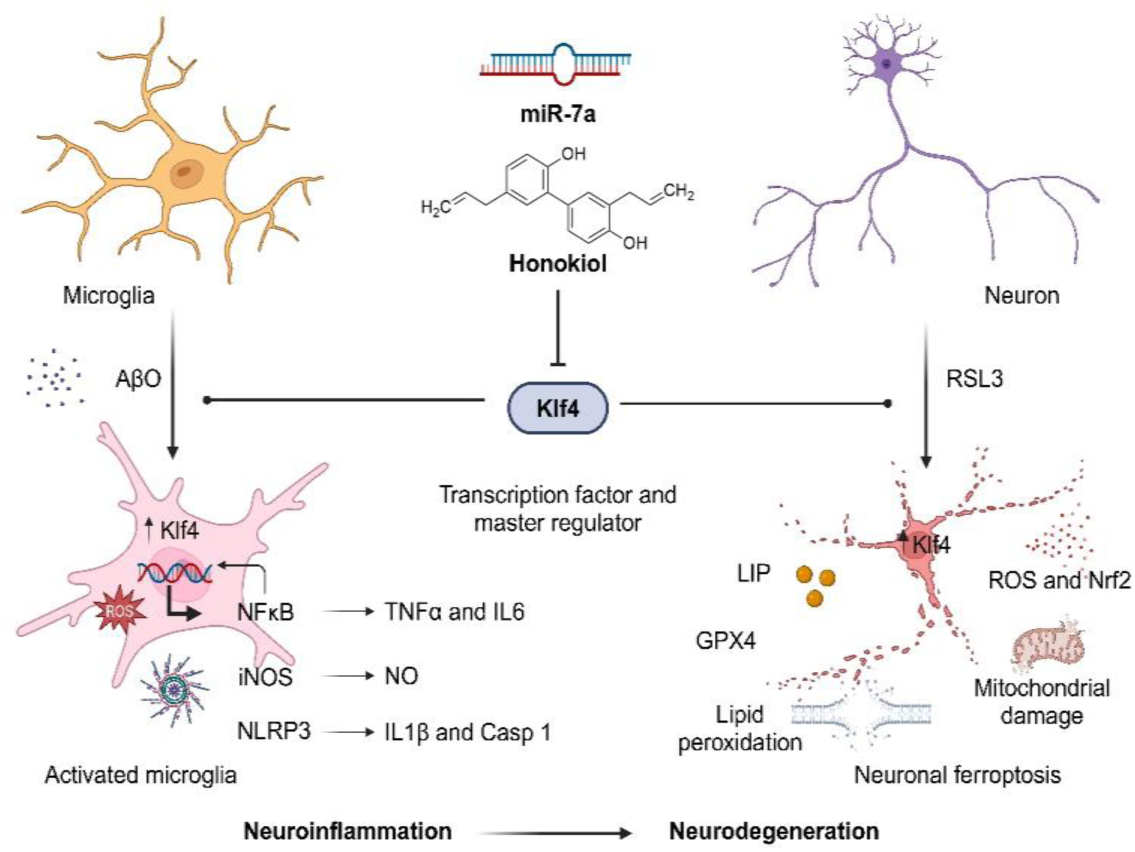

## INTRODUCTION

AD is a prevalent progressive neurodegenerative condition, characterized by the accumulation of amyloid-beta (Aβ) and tau protein aggregation species^1,2^. These pathological events are responsible for the observed impairments in learning, memory, and overall cognitive function [1, 2]. The multifaceted etiology of AD, with its intricate pathological indicators and clinical symptoms, presents formidable challenges in the development of effective treatments [3]. Currently available therapeutic measures are confined to providing only temporary and symptomatic relief [4]. The stagnation in therapeutic advancements can be largely attributed to the insufficient exploration of viable therapeutic targets and a dearth of innovative strategies aimed at accurately targeting the regulators of disease pathology [5]. There is an urgent and imperative need to identify new therapeutic targets that are intrinsic to the neurodegenerative processes, to improve the therapeutic management of AD.

Recent studies have expanded the understanding of AD pathogenesis beyond Aβ and tau to include the pivotal role of neuroinflammation in the disease onset and progression [6]. Microglial cells, the brain resident macrophages, serve a paradoxical role. While microglia are essential for the function and homeostasis of central nervous system (CNS), they also exacerbate neurodegeneration by releasing proinflammatory mediators [7]. Microglia exist in homeostatic and disease-associated states that regulates its dual role [8, 9]. However, the molecular mechanisms governing these states remain elusive. Unravelling the key regulators of microglial state and neuroinflammation is crucial for developing therapeutic interventions [10, 11]. Activated microglia drives neuroinflammation via proinflammatory mediators, modulated by pathways such as NF-κB, reactive oxygen species (ROS), and NLRP3 inflammasome [12]. Ferroptosis/oxytosis, a new form of cell death triggered by iron mediated lipid peroxidation. It is characterized by elevated labile iron pool (LIP), lipid peroxidation, reactive oxygen species (ROS) production, oxidative stress, impaired GPX4 function, disrupted Nrf2 signalling, mitochondrial damage, and subsequent cell death implicated in various neurodegenerative disorders [13–18]. Recent studies have established the connection between ferroptosis and various neurodegenerative diseases, including AD [14, 19]. Proteins involved in the ferroptosis pathway are significantly altered in AD, implicating its potential contribution to neurodegeneration [20–22]. The increased labile iron levels and lipid peroxidation products in the AD brain suggest its pathological nexus with ferroptosis [14, 23]. Evidence of ferroptosis in aging human microglia and Aβ-induced ferroptotic signatures in neuronal cells underscore its significance in cell death mechanisms [14, 23, 24]. A deeper understanding of the molecular underpinnings and regulators of ferroptosis may offer new avenues for mitigating neurodegeneration in AD.

MicroRNAs (miRNAs) play functional and regulatory roles in cellular physiology and pathophysiology. They are the master regulators of gene expression at the posttranscriptional level, orchestrating the degradation of messenger RNAs (mRNAs) or suppressing their translation [25]. As master regulators, miRNAs modulate a multitude of mRNA targets, influencing physiological and pathological pathways [26–29]. They orchestrate various neuronal functions, from signal transduction to memory consolidation. Despite their significance and critical role, comprehensive identification and understanding of miRNAs in AD pathology are still limited. MiRNAs have been shown to target APP, BACE1, AChE, tau, GSK3β, and phosphatases [28]. These insights underscore the potential of miRNA and miRNA-based therapeutics in targeting AD pathological mechanisms. Discovering novel miRNAs that govern other pathways contributing to neurodegeneration, such as neuroinflammation and ferroptosis, herald a new therapeutic prospect.

In this study, we embarked on a comprehensive analysis of miRNA profiles and their target mRNAs to establish a miRNA-mRNA-pathway network employing the APP/PSEN1 transgenic (Tg) AD mouse model. The study uncovered a significant upregulation of miR-7a in the AD brains, highlighting its regulator function in AD pathology. Notably, *Klf4* was identified as a principal mRNA target of miR-7a and Klf4 protein levels was significantly elevated in AD. Klf4 was upregulated in AβO induced neuroinflammation and miR-7a-Klf4 axis was found to regulate microglial activation and neuroinflammation. Klf4 is a pivotal factor in neuronal ferroptosis and regulate ferroptotic neuronal death. The inhibition of Klf4 by miR-7a effectively reduced Ab oligomer (AβO)-induced microglial activation, neuroinflammation, and RSL3-triggered ferroptotic cell death. The pharmacological modulation of the miR7a-Klf4 axis by honokiol is a promising approach to alleviate neuroinflammation and ferroptosis in AD, offering potential RNA-based and small molecule interventions.

## MATERIAL AND METHODS

### Chemicals and reagents

All the chemicals and reagents were purchased and used without purification unless mentioned. The RNA isolation reagent Trizol, DEPC treated water (#AM9922), nuclease free water (NFW) (#AM9932), RNaseZap, Verso cDNA Synthesis Kit (#AB1453A), DyNAmo ColorFlash SYBR Green qPCR Kit (#F416L), Lipofectamine™ RNAiMAX Transfection Reagent (#13778075) and MicroAmp™ Optical 96-Well Reaction Plate with Barcode & Optical Adhesive Films (#4314320), Calcein AM (# C3099), DAPI (#62248), BODIPY™ 581/591 C11 (Lipid Peroxidation Sensor) (#D3861) MitoTracker™ Green FM Dye (# M46750) Rhodamine 123 (# R302), Human beta Amyloid (1-42) PTD Recombinant Protein (# 03-112) were purchased from Invitrogen, ThermoFisher Scientific. miRNA mimic, inhibitor (Assay Id - MH10047) and Negative Control (# 4464058) are purchased from Invitrogen, ThermoFisher Scientific and Eurogenetec. iTaq Universal SYBR Green Supermix 2X (#1725121) was purchased from BioRad. miScript II RT Kit (#218161) was purchased from QIAGEN Pvt Ltd. The custom DNA oligos (primers) were synthesized by Eurofins and directly used without further purification. Atto-647 5’ modified miR-7a RNA sequence and Klf4 mRNA fragment RNA oligonucleotides were purchased from IDT (Supplementary Table S10). 2′,7′- Dichlorofluorescin diacetate (DCFDA #287810), Griess reagent (#G4410), 1S,3R-RSL 3 (#SML2234), *E. coli* lipopolysaccharide (LPS) (#L4391), 3-(4,5-dimethylthiazol-2-yl)-2,5 diphenyl tetrazolium bromide (MMT) (#M5655), Quercetin (#Q4951), honokiol (#H4914) and nicorandil (#N3539) from Merck, Sigma Aldrich. The cell media Dulbecco’s Modified Eagle Medium (DMEM-F12), fetal bovine serum (FBS), goat serum (GS), penicillin-streptomycin (PS), Roswell Park Memorial Institute (RPMI) were purchased from Gibco, ThermoFischer Scientific. Sterile T25 flasks, 96. 12 and 6-well plates (Thermos Fischer Scientific), and confocal dishes (SPL life Sciences) were purchased and used without further sterilization. The sandwich ELISA kits for mouse IL1β (#RK00006), TNFα (#RK00027) and IL6 (#RK00008) were purchased from ABclonal and used as per the manufactures protocol. The cDNA synthesis was performed on Eppendorf thermocycler and qRT-PCR was performed on QuantaStudio 5, Applied Biosystems real time PCR instrument. Circular dichroism (CD) and fluorescence spectra were recorded on J1500 (Jasco) and Agilent Cary Eclipse Fluorescence Spectrophotometer respectively. Well plate absorbance and fluorescence measurement were performed in Spectramax i3 (Molecular devices) plate reader instrument. Live cell fluorescence imaging was carried out in Leica DMi8 fluorescence microscope equipped with live cell set up. The confocal imaging was performed using Olympus Fluoview-3000 confocal laser scanning microscope and analyzed by ImageJ software. The list of antibodies and their details used in the study are detailed in supplementary table S1.

### Animals and maintenance

APP/PSEN1 double transgenic mice [B6.Cg-Tg(APPswe,PSEN1dE9)85Dbo/Mmjax] were obtained from the Jackson Laboratory (MMRRC stock no 34832) and maintained in JNCAR Animal Facility under 12 h light/dark cycle. All the animal breeding, maintenances and studies were performed according to the protocols of the Institutional Animal Ethics Committee (IAEC). The protocol (TG004) was approved by the IAEC.

### RNA extraction and NanoString miRNA expression assay

Total RNA including miRNA was isolated from mouse cortex and hippocampus tissue using Qiagen miRNAeasy mini kit (Cat#217004). Tissue was homogenized in TOMY homogenizer in presence of liquid nitrogen, for three cycles. 700 µl of QIAzol Lysis reagent was added to the tissue powder and mixed well. Lysate was incubated at room temperature for 5 minutes and 140 µl of chloroform was added for phase separation. Rest of the protocol was followed as per manufacturer’s guidelines. RNA was eluted in 25µl of nuclease free water (Ambion, Cat #AM9932). Quantitation was performed using Qubit RNA HS assay (Invitrogen, Cat #Q32855) kit and qualitatively analyzed on Agilent 2100 bioanalyzer nano chip (Agilent, Cat #5067- 1511). With NanoString Mouse v1.5 -miRNA assay kit (CSO-MMIR15-12), miRNA (3μl) was ligated to mir-Tag with ligation buffer and ligase supplied with the kit. Ligated product was diluted with 15 µl of nuclease free water and denatured at 85 °C for 5 minutes. 5 µl of this was hybridized overnight at 65 °C with reporter and capture probes. Protocol followed as per nCounter miRNA expression assay user manual (MAN-C0009-07). Post hybridization, samples were analyzed on NanoString nCounter SPRINT machine.

### NanoString nCounter miRNA profiling and data analysis

Analysis was performed on all samples using the nCounter Analysis System (NanoString Technologies) and the nCounter Mouse v1 miRNA Panel (NS_M_miR_v1.5) having 599 unique miRNA barcodes for endogenous miRNA. The housekeeping genes used in the panel includes, beta-actin (Actb), beta-2-microglobulin (B2m), glyceraldehyde 3-phosphate dehydrogenase (Gapdh) and ribosomal protein L19 (Rpl19). The panel also includes the SpikeIn mIRNAs, *Arabidopsis thaliana* miR159a (ath-miR159a), *Caenorhabditis elegans* (cel)- miR-248 and miR254, and *Oryza sativa* (osa)-miR 414 along with the positive and negative controls for assessing overall assay efficiency as well as for monitoring the ligation efficiency.

The raw miRNA data in.RCC (Reporter Code Count) format was further analyzed using nSolver analysis software (NanoString technologies) version 4.0. QC metrics of imaging, binding density, positive control, limit of detection and ligation were checked before proceeding with further analysis. Normalization of raw data was performed using the geometric mean of positive controls and top 100 highly expressed miRNAs. The differential expression between the groups were calculated using the build ratio utility present within the nSolver (foldchange), which uses Student’s unpaired t-test for the *p* value calculation. The thresholds considered for a significant differentially expressed miRNA were, foldchange ≥ 1.2 or ≤ -1.2, P-value ≤ 0.05 and either of the two groups (test/control) should have their geometric mean expression ≥ average count for negative control probes in the panel. Unsupervised hierarchical clustering analysis followed by heatmap visualization and principal component analysis (PCA) was performed with the normalized and log2 transformed data using ClustVis (https://biit.cs.ut.ee/clustvis/) [30].

### MiRNAs target prediction and functional over-representation analysis

For the statistically significant miRNA features (*P* value < 0.05), mRNA target prediction was performed in a dual approach of experimentally validated mRNA target prediction and *in-silico* target prediction. Experimentally validated targets were predicted using miRTarBase v.9.0 database, miRecords v.4 and TarBase v.8. The union of all interactions retrieved from the 3 queried sources was considered as the set of validated miRNA–target gene interactions in our study. To search for additional miRNA targets, *in-silico* target prediction were performed for selected miRNAs using TargetScan v.8 and miRDB v.6.0 queried using miRWalk webserver, where the score for target interaction probability threshold was set to 0.9. Targets where the miRNA binding sites ranged from cds, 5’- and/or 3’-UTR regions were considered for analysis. Only the intersection of all interactions retrieved from the 2 queried *in-silico* sources were considered as the set of *in-silico* predicted miRNA–target gene interactions in our study. Further the Gene Ontology (GO) Biological Process, Cellular Component and Molecular Function along with REACTOME pathways were queried for both the targets of up or downregulated miRNAs, separately using ClusterProfiler [31] and ReactomePA R package [32]. Functions/Pathways with a *p*-value <0.05 were considered strongly enriched for the target genes targeted by selected miRNAs.

### MiRNA-mRNA-pathway network analysis

Validated target collection of selected individual miRNAs (*p* value < 0.05) were enriched for Reactome Pathways using DAVID webserver (https://david.ncifcrf.gov/) [33]. The top significant (p value < 0.05) enriched pathways and their corresponding genes were selected for further visualization as miRNA-mRNA-Pathway network using Cytoscape v3.10 [34]. The miRNA node size has been scaled based on the total number of validated interactions shared with target mRNAs.

### MiRNA disease association and miRNA-mRNA-disease network

In order to associate or enrich the differentially regulated miRNAs between AD relative to healthy controls (*p* value < 0.05), both the association of miRNA with disease as well as validated targets of miRNA with disease were analyzed. The miRNAs and disease association was established using PhenomiR v.2.0 and miEAA (querying MNDR database), while the miRNA-validated target and disease association was analyzed through MNDR v.3.0 and DisGeNET cytoscape application [35]. Diseases with a *p*-value <0.05 were considered strongly enriched for the selected collection of miRNAs. The miRNA-Gene-Disease network was visualized using Cytoscape v3.10.

### MiRNA expression validation by qPCR assay

The total RNA used for NanoString was further utilized for validation by qPCR. The miRNA cDNA was synthesized by miScript II RT Kit by using manufactures protocol. The primers for the target miRNA were designed by using miRPrimer software [36] and custom synthesized from Eurofins. The assay were optimized and the qPCR was performed in Applied Biosystems QuantaStudio 5 system and assessed the miRNA expression with snoRNA as control with comparative Ct (ΔΔCT) method.

### Immunofluorescence of mouse brain

The brain tissue was fixed in a 4% paraformaldehyde solution, then dehydrated in a 25% sucrose solution. The tissues were sectioned using a cryotome with a thickness of 25 μm. The sections were washed twice with PBS to remove the antifreeze media and permeabilized with PBS containing 0.01% Triton X-100 for 10 minutes. After two additional washes with PBS, the tissues were incubated with a blocking solution (PBST with 3% goat serum). The tissues were then treated with the Klf4 primary antibody overnight at 4 °C, washed three times with PBS, and subsequently treated with an Alexa-488 fluorophore-conjugated secondary antibody at room temperature. After a 1-hour incubation, the samples were washed three times with PBS, mounted with DAPI-containing mounting media for nuclear staining, and subjected to confocal microscopy,

### Western blot

Twelve-month-old WT and AD mice were sacrificed, and brain samples were collected. The brain tissues were homogenized in RIPA lysis buffer containing a protease inhibitor cocktail using a Dounce homogenizer. After complete homogenization, the lysate was centrifuged at 12,000 rpm for 10 minutes at 4 °C, and the supernatant was collected. Total protein concentration was quantified using the Bradford assay with BSA as a standard. Forty micrograms of total protein were subjected to SDS-PAGE followed by western blotting. The blots were blocked with 5% skim milk powder, incubated with Klf4 primary antibody (1:1000) overnight at 4 °C, and then treated with HRP-conjugated secondary antibody (1:5000) for 1 hour at room temperature. The blots were developed on X-ray films using Clarity Western Enhanced Chemiluminescence (ECL) reagent (BIORAD). Band intensities were quantified using Fiji.

### Circular dichroism (CD) and fluorescence spectroscopy

To understand the binding of miR-7a with the 3’-UTR region of Klf4 mRNA, we performed CD spectroscopy. Custom-synthesized miR-7a and the complementary region of Klf4 mRNA sequences were purchased and dissolved in Tris HCl buffer (10 mM, pH 8.0). The miR-7a (25 μM) and Klf4 sequences (25 μM) were annealed at 65 °C in 10 mM tris buffer, and their CD spectra were measured individually. The sequences were then mixed, annealed at 65 °C to promote binding at the complementary region, and allowed to cool slowly. The mixture was subjected to CD spectroscopy, and the data were processed and plotted using Origin software. To further determine the binding ability of miR-7a with Klf4, fluorescence-tagged miR-7a (miR-7a-Atto647) was titrated with increasing concentrations of the Klf4 sequence. A fixed concentration of miR-7a-Atto647 (500 nM) was used, and Klf4 sequence was added incrementally (250 nM to 10 μM), with Atto647 fluorescence measured after a 5-minute incubation period following each addition. The complementary base pairing between miR-7a and Klf4 resulted in changes in Atto647 fluorescence, attributed to alterations in the local environment of the fluorescent dye due to base stacking. The change in fluorescence was plotted against the concentration of the Klf4 sequence to assess the binding behavior of miR-7a and Klf4.

### General cell culture

SH-SY5Y neuroblastoma cells were cultured in DMEM-F12 medium supplemented with 10% FBS and 1% PS and incubated in a humidified incubator at 37 °C with 5% CO_2_. The cells were grown in T25 flasks, trypsinized at 80% confluency, and reseeded as needed. N9 microglial cells were cultured in RPMI medium with 10% FBS and 1% PS and incubated under the same conditions. For experiments, 96-well, 12-well, and 6-well plates were seeded with cell densities of 15,000, 80,000, and 200,000 cells per well, respectively, and cultured for 24 hours before proceeding with experiments.

### RT-PCR and immunofluorescence

MiR-7a, Klf4 mRNA, and protein expression in SH-SY5Y and N9 microglial cells were assessed using qPCR and immunofluorescence. For qPCR, cells were cultured in a 6-well plate under standard conditions. Total RNA was isolated using Trizol reagent and quantified using a nano spectrophotometer to assess quantity and quality. For miR-7a expression analysis, cDNA was synthesized using the miScript II RT Kit according to the manufacturer’s protocol. Expression levels were analyzed by qPCR with snoRNA as the housekeeping gene, and the ratio of Ct values was plotted to compare expression between the two cell types. Similarly, for *Klf4* mRNA expression, cDNA was prepared using the Verso cDNA Synthesis Kit, and mRNA expression was assessed by qPCR using β-actin as the control gene. For Klf4 protein expression, cells were cultured in confocal dishes for 24 hours and then subjected to immunofluorescence. Cells were washed twice with PBS for 5 minutes each, fixed with 4% paraformaldehyde for 15 minutes, and washed three times with PBS for 5 minutes each. The cells were then permeabilized for 5 minutes, washed twice with PBS, and blocked for 30 minutes. Cells were treated with the Klf4 primary antibody (1:100) overnight at 4 °C, washed three times with PBS for 5 minutes each, and then incubated with the Alex-488 fluorophore-conjugated secondary antibody (1:500) for 1 hour at room temperature. After washing twice with PBS for 5 minutes each, DAPI solution in PBS was added for 15 minutes to stain the nuclei, and unbound probe was washed away with PBS. Finally, the samples were subjected to confocal imaging.

### MiR-7a mimic and inhibitor transfection and immunofluorescence experiments

The miR-7a mimic, inhibitor and negative control (NC) were purchased from Invitrogen and Eurogentec used as per the manufacturers protocol. The products were constituted in nuclease free water and used for the experiments. For transfection the cells were cultured in confocal dishes and transfected with miR-7a mimics and inhibitor using Lipofectamine™ RNAiMAX transfection reagent as per the standard procedure. Cells were incubated for 9 h and then added low serum media and after 30 h further subjected immunofluorescence for protein expression.

### Inflammation experiments - qPCR and immunofluorescence

N9 microglial cells were seeded in confocal dishes and 12-well plates and cultured under standard conditions for 24 h. The cells were treated with different concentrations of LPS (100, 250, and 500 ng/ml) and incubated for an additional 24 hours. Total RNA was isolated and subjected to qPCR for mRNA expression analysis. Cells in the confocal dishes were fixed, subjected to Klf4 immunofluorescence, and imaged using a confocal microscope. For AβO-induced inflammation, cells were primed with LPS (500 ng/ml) for 4 hours, then treated with AβO oligomers (2.5, 5, and 10 μM). After treatment, the cells were subjected to qPCR for mRNA expression analysis and Klf4 immunofluorescence for protein expression.

To understand the inhibitory effect of microglial activation by miR-7a mimics, the microglial cells were transfected with miR-7a mimic, after 9 h the media was replaced with low serum media. The cells were primed with LPS (500 ng/ml), followed by treatment with AβO (5 μM) for 24. Then cells were subjected to Iba1 immunofluorescence to analyses the inhibitory effect of miR-7a on microglial activation.

### DCFDA assay for ROS measurement

To assess ROS generation and the rescuing effect of miR-7a, microglial cells were transfected with miR-7a mimic (100 and 250 nM). After 9 hours, the medium was replaced with low serum medium. The cells were primed with LPS (500 ng/ml), followed by treatment with AβO (5 μM) for 24 hours. Subsequently, the cells were subjected to the DCFDA assay to measure intracellular ROS. Cells were washed twice with PBS, then treated with 20 μM DCFDA reagent in serum-free RPMI medium. After 45-minute incubation, the cells were washed twice with PBS, placed in fresh medium, and analyzed using a multiplate reader in well scan mode to measure the mean fluorescence (λ_ex_ = 485 and λ_ex_ = 525) at from each well. A minimum of 100-point readings was set in a circular pattern, and the mean DCF fluorescence was used to determine cellular ROS levels.

### Griess assay for NO measurement

The microglial cells were transfected with miR-7a mimic, after 9 h the media was replaced with low serum and phenol red free media. The cells were primed with LPS (500 ng/ml) for 4 h, followed by treatment with AβO (5 μM) for 24 h. Cell media was subjected to Griess assay to detect the relative NO released by activated microglial cells. Equal volume of cell media and Griess reagent (100 μL) were mixed and incubated for 10 min. The absorbance of the chromophore formed during diazotization of nitrite with sulphanilamide and subsequent coupling with naphthylethylenediamine was recorded by absorbance at 546 nm. The relative NO levels were calculated by normalizing against control cell media.

### Neuroinflammatory mRNA and protein expression

To understand the inhibitory effect of microglial activation by miR-7a mimics, the microglial cells were transfected with miR-7a mimic, after 9 h the media was replaced with low serum media. The cells were primed with LPS (500 ng/ml), followed by treatment with AβO (5 μM) for 24. Then cells were subjected to qPCR for mRNA expression quantification of various neuroinflammatory mediators and immunofluorescence for protein expression analysis. The cell media was subjected to sandwich ELISA for protein quantification of TNFα, IL6 and IL1β using Abclonal ELISA kit as per the manufacturers protocol.

### Ferroptosis experiments - qPCR and immunofluorescence

SH-SY5Y cells were cultured in 12 well plate and confocal dishes and treated with different concentrations of RSL3 (0.5, 1 and 2.5 μM) and vehicle to control at 60% confluency and incubated for 24 h. Then the cells were subjected to qPCR and immunofluorescence of Klf4 to understand the Klf4 levels in RSL3 induced ferroptosis condition.

Cells were transfected with miR-7a mimic at 50% confluency, after 9 h the media was replaced with low serum media. After 6 h incubation the cells were treated with RSL3 (2 μM) and incubated for 24 h. The samples were subjected to GPX4 and Nrf2 immunofluorescence. Mean fluorescence intensity was calculated for relative protein levels and the ratio of nuclear and cytoplasmic mean fluorescence used to measure the extent of nuclear translocation of Nrf2.

### Lipid peroxidation assay – Bodipy sensor

The extent of lipid peroxidation in the cells was imaged using the Bodipy C11 fluorescent probe. The probe shows red emission and upon reacting with lipid peroxides display green fluorescence. The increase in green fluorescence shows increased lipid peroxidation. The cells were cultured and transfected with miR-7a mimic and then treated with RSL3 (2 μM) and incubated for 24 h. Then cells were treated with Bodipy sensor (5 μM) and incubated for 30 min followed by washing with PBS and imaging in live cell set up. The induction of ferroptosis is marked by increased lipid peroxides and the increased green fluorescence and the rescue has reduced green emission. To quantify the extent of lipid peroxidation we have performed assay in 24 well plate. Fluorescence in green (λ_ex_ = 488 and _λem_ = 510 nm) and red (λ_ex_ = 581 and _λem_ = 591 nm) emission were recorded in plate reader with well scan mode and plotted ratio of green and red to calculate relative lipid peroxidation.

### Labile iron pool (LIP) measurements – Calcein-AM imaging

Cells were seeded in confocal dishes and cultured in standard conditions. At 50% confluency the cells were transfected with miR-7a mimic using RNAi max transfection reagent and then after 9 h the media was replaced with low serum media. After 12 h the cells were treated with RSL3 (2 μM) to induce ferroptosis for 24 h. Then cells were treated with calcein-AM (500 nM) for 45 min and washed with PBS and imaged in live cell imaging system. Calcein-AM intracellularly binds to Fe^2+^ and exhibits reduced fluorescence indicating increased labile iron pool. The cells were imaged in green channel with live cell set up. The mean fluorescence intensity was quantified using Image J Fiji and normalized against control for relative iron levels in cells.

### Mitochondrial imaging and MMP measurement

To analyze the protective effect of miR-7a targeting Klf4 on the ferroptosis mediated mitochondrial damage we performed mitochondrial imaging and estimation of mitochondrial membrane potential (MMP). To image the mitochondrial damage the cells transfected with miR-7a (100 nM and 250 nM) and challenged with RSL3 were incubated for 24 h. then the cells were treated with Mitotracker green GM (250 nM) and incubated for 30 min. The cells were washed twice with PBS and imaged in fluorescence microscope with green channel in live cell set up. To measure the MMP cells cultured in 24 well plate were transfected with miR-7a (50 nM, 100 nM and 250 nM) and then challenged with RSL3 and incubated for 24 h. Cells were treated with Rho123 (250 nM) a potential dependent mitochondrial probe and incubated for 30 min. Washed with PBS and fluorescence (λ_ex_ = 507 and λ_em_ = 527 nm) was measured in multiplate reader with well scan mode with 80 reading points per well and the mean fluorescence intensity was used to measure the MMP.

### Cell rescue by miR-7a and compounds

SH-SY5Y cells were seeded and cultured in 96 well plates and transfected with miR-7a mimic using RNAi transfection reagent. After 6 h media was removed and incubated with low serum DMEMF12 media. 6 h later the cells were treated with 2 μM of RSL3 to induce the ferroptosis. After 30 h of incubation RSL3 induced ferroptotic cell death and rescue by miR-7a was assessed by MTT assay. Briefly cells were treated with 10 μl MTT reagent (5mg/mL) for 3 h. Then the formazan crystals were dissolved in DMSO:Methanol (1:1) solvent mixture. The absorbance was measured at 570 nm to calculate the percentage cell viability against the control with vehicle treatment.

The cells were seeded and cultured in 96 well plates for 24 h and at 70 % confluency the cells were treated with AβO (5 μM) for N9 microglial cells for inflammation or RSL3 (2 μM) for ferroptosis in SH-SY5Y cells and with different concentration of compounds. After 30 h of incubation the cell viability was assessed by MTT assay. Percentage cell viability was calculated with respect to control using the absorbance at 570 nm.

### RT-PCR and immunofluorescence with honokiol treatment

SH-SY5Y and N9 cells were cultured in confocal dishes and at 60 % confluency the cells were treated with AβO alone and with honokiol (2.5 and 5 μM) to induce microglial activation and ferroptosis. After 24 h incubation the cells were subjected to Iba1 immunofluorescence. The microglial activation and rescue by honokiol was assessed using Iba1 positivity and rescue by honokiol treatment. The effect of honokiol on *Klf4* expression was assessed by RT-PCR and immunofluorescence. SH-SY5Y cells were cultured in 12 well plate and confocal dishes and then treated with honokiol (2.5 μM). Total RNA was isolated from well plate and subjected to RT-PCR for *Klf4* mRNA expression. The dishes were subjected to immunofluorescence and the Mean fluorescence intensity of Klf4 was calculated for relative protein levels.

### Mouse cortical primary-glia mixed culture

To evaluate anti-inflammatory and neuronal rescue effect of honokiol we have employed mouse primary cortical neuron-glia mixed culture using reported protocol with slight modifications [37]. Day one pups (P1) were sacrificed and dissected to get the cortical tissue and subjected to trituration. The tissue was triturated sequentially with serological pipette, 1 ml, 200 μl and 20 μl tips to get homogeneous cell suspension. Then the cells were strained through 100 μm mesh size filter and cells were pelleted by centrifugation. The cells were seeded onto poly-D-lysine coated (20 μg/ml) dishes and plates. Then the cells were maintained in neuron-glia maintenance media for 8 days. Cells were treated with AβO (2.5 μM) for 24 h and subjected to cell viability and immunofluorescence assay. Neuronal cells and activated microglia were probed with NeuN (Rabbit) and Iba1 (Chick) primary antibody (1:200 dilution) respectively. Alex-488 conjugated anti-rabbit and Alex-568 conjugated anti-chicken secondary antibodies (1:500 dilution) were used and subjected to confocal imaging.

### Statistical analysis

All statistical analysis was performed by using R (version 4.0.2, https://www.r-project.org/) for transcriptomic analysis. P value below 0.05 was considered statistically significant. Pairwise comparisons were used to compare between the WT and AD groups. We have used GraphPad and Origin for plotting and analysis of the data. The statistical tests were applied using GraphPad. Used student t test for comparing two groups with * is p>0.005 and One Way ANOVA to compare multiple groups with Bonferroni Post Hoc test * is p>0.005 and ns indicate no significance difference.

## RESULTS

### Differentially expressed miRNAs and altered miR-mRNA-pathway/disease network associated with AD

A multitude of miRNAs are dysregulated in AD, and miRNA panels have been widely utilized as diagnostic biomarkers. The extensive regulatory roles of miRNAs within the CNS have prompted an in-depth investigation into their pathogenic and neuroprotective implications in AD. Utilizing the APP/PSEN1 transgenic mouse model, we assessed the expression of miRNAs in AD-afflicted brains in comparison to wild type (WT) mice, aiming to establish altered miRNAs, mRNAs, and pathway network (Fig. 1A). The transcriptomic profiling of miRNAs in the cortex and hippocampus regions was carried out using the NanoString platform (Supplementary Table S2 and S3). Transcriptome analysis revealed that 21 out of 577 analyzed miRNAs were differentially expressed (nCounter counts threshold of 40 and p<0.005, t-test) in the cortex of AD compared to WT brain (Fig. 1B and D). Notably, 6 miRNAs (miR-7a, miR-696, miR-34a, miR-302b, miR-151-3p) exhibits more than 1.5-fold change in expression levels. MiR-7a being significantly upregulated with a fold change of 2.05, whereas miR-696, showed pronounced downregulation by 2.55-fold. In the hippocampus, 11 miRNAs showed substantial expression changes, with a mix of both upregulation (5) and downregulation (6) observed (Fig. 1C and S1C). The analysis of 577 miRNAs highlighted consistent upregulation of miR-7a in both cortex and hippocampus, while miR-148 and miR-301a were specifically upregulated in the hippocampus. Conversely, miR-696 and miR-34a were notably downregulated in the cortex, with miR-153 and miR-128 downregulated in the hippocampus. Multidimensional plots distinctly illustrated the separation of WT and AD miRNA profiles within both brain regions, indicating the potential regulatory impact of these dysregulated miRNAs in AD pathology (Fig. S1A and B).

**Figure 1.**
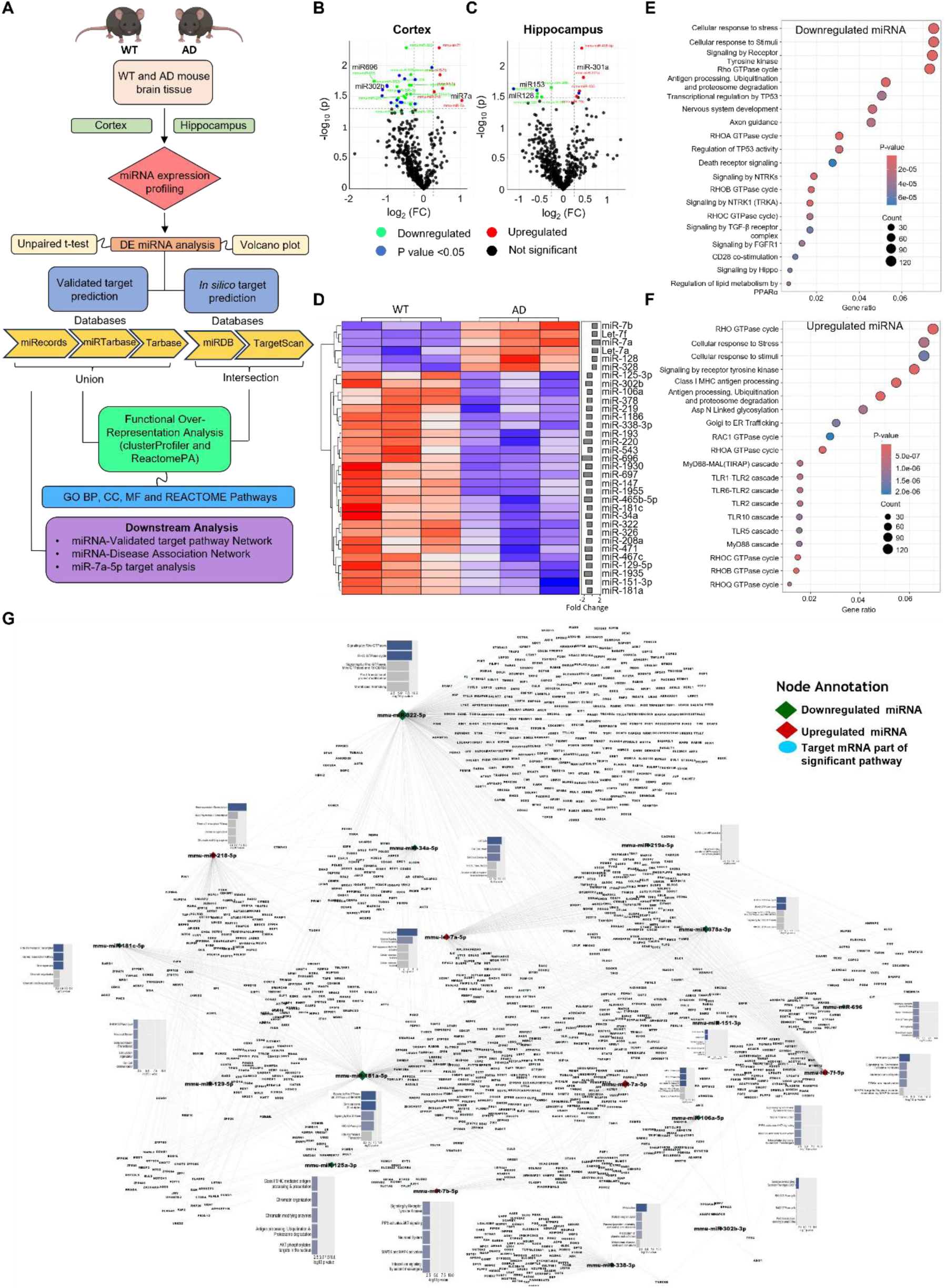
Differentially expressed miRNAs and altered miRNA-mRNA-pathway/disease network. (**A**) Schematic of experimental and bioinformatics design of miRNA transcriptomics analysis. Volcano plots showing differentially expressed miRNAs in cortex (**B**) and hippocampus (**C**) of 12-month-old APP/PSEN1 Tg mouse model. Differentially expressed miRNAs were identified using NanoString analysis (n = 3 mice per group, P < 0.05). (**D**) Heatmap of the top variable miRNAs of cortex. Molecular pathway of functional overrepresentation analysis for identified mRNA targets of upregulated (**E**) and downregulated (**F**) miRNA in cortex (*p*-value <0.05). (**G**) MiRNA-mRNA-pathway network constructed for dysregulated miRNA of cortex and hippocampus by using significantly (p value < 0.05) enriched pathways and their corresponding genes.

Next, we performed a functional over-representation analysis on the altered mRNAs, employing target mRNA predictions for dysregulated miRNAs (Supplementary Table S3 and 4). Further, we interrogated the Gene Ontology (GO) categories - Biological Process, Cellular Component, and Molecular Function and REACTOME pathways for targets of both upregulated and downregulated miRNAs, utilizing ClusterProfiler and ReactomePA tools. The functional over-representation analysis of downregulated miRNAs in the cortex, using predicted mRNAs, identified pathways associated with AD pathology, including cellular stress response, transcriptional regulation, nervous system development, antigen processing, death receptor signaling, cell cycle signaling, TGF-β signaling, and lipid metabolism (Fig. 1E and S2). Analysis of upregulated miRNA targets indicated involvement in stress response, antigen processing, proteasomal degradation, cargo trafficking, immune cell signaling, toll-like receptor cascade, and Rho GTPase cycle pathways (Fig. 1F and S3). In the hippocampus, downregulated miRNA pathways encompassed Rho GTPase cycle, cytokine signaling, antigen processing, proteasomal degradation, nervous system development, Golgi trafficking, transcriptional regulation, SUMOylation, and lipid metabolism (Fig. S4). The pathways for upregulated miRNAs included Rho GTPase, transcriptional regulation, TLR cascades, MAPK activation, and signaling by the Hippo pathway (Fig. S5). Collectively, the functional over-representation analyses elucidated that dysregulated miRNA target mRNAs involved in various neuronal and microglial related pathways, which are potentially implicated in the development and progression of the disease. This underscores the critical role of miRNAs in deciphering the intricate mechanisms of neurodegenerative diseases such as AD.

The validated mRNA targets of selected individual miRNAs (p value < 0.05) were enriched for reactome pathways using DAVID webserver. The most significantly enriched pathways and their corresponding genes were subsequently visualized within a miRNA-mRNA-Pathway network. This network demonstrated the regulatory functions of miRNAs across various pathways, including transcription regulation, chromatin organization, Rho GTPase cycle, post-translational modifications (PTM), membrane trafficking, cell-cell communication, immune cell signaling, cellular stress response, cell cycle pathways, AMPA receptor activity, and synaptic plasticity (Fig. 1G, S6A and Supplementary Data 1 and 2). These pathways have known associations with AD and other neurodegenerative diseases. Further investigations were undertaken to delineate the connections between miRNAs, validated mRNAs, and diseases, resulting in the construction of a miRNA-mRNA-disease network. The relationships between miRNAs and diseases were established using PhenomiR v.2.0 and miEAA (querying the MNDR database). Further, the associations between miRNA-mRNA and diseases were analyzed using MNDR v.3.0 and the DisGeNET Cytoscape (Supplementary Table S6-9). The miRNA-mRNA-disease network distinctly illustrated the links between dysregulated miRNAs and various neurodegenerative disorders, including AD (Fig. S6B and C). These comprehensive transcriptomic and bioinformatic analyses have solidified the association of identified miRNAs with pathological pathways implicated in AD and other neurodegenerative diseases, emphasizing their potential as diagnostic and therapeutic targets.

### MiR-7a is significantly induced and directly target Klf4 in AD

After identifying the altered miRNAs in the cortex and hippocampus of AD brains and the establishment of their mRNA-pathway-disease network, we validated significantly altered miRNAs (miR-7a, miR-148a, miR-34a, miR-128a, and miR-153a) expression in the AD cortex and hippocampus using qRT-PCR. The expression patterns aligned with our comprehensive transcriptome analysis, revealing a significant upregulation of miR-7a and miR-148, while miR-34a and miR-128a were notably downregulated (Fig. 2A). Among these, miR7a was markedly upregulated in both brain regions. Considering the neuroprotective role of miR-7a suggested by various studies, we hypothesize that miR-7a is potentially induced under disease conditions to mitigate AD pathology. The pathway network analysis vide supra revealed altered transcription regulation, immune, cell stress and cell death response pathways. Our literature analysis, target prediction, and sequence alignment suggest *Klf4* as a target for miR-7a bearing potential role in AD pathology. We observed an increase in Klf4 levels in the AD brain by immunofluorescence and western blot, suggesting its potential role in AD onset and progression (Fig. 2B, C and Supplementary Data 3). The conserved sequence of miR-7a aligns complementarily with the 3’ UTR region of *Klf4* mRNA, indicating a potential binding interaction that is reported to attenuate Klf4 expression (Fig. 2D). To investigate the complementary base pairing between miR-7a and Klf4, we utilized circular dichroism (CD) spectroscopy, which differentiates between single-stranded and double-stranded RNA structures. The interaction between miR-7a and *Klf4* resulted in a partial double-stranded RNA formation, as evidenced by characteristic CD spectral bands at 270 (+), 240 (–), 220 (+), and 210 (–) nm (Fig. 2E). The annealing of miR-7a with the complementary Klf4 fragment led to a shift in the CD spectral band, indicative of base pairing which resulted in shifting of 270 nm band to lower wavelength and lowered intensity at 210 (–) nm with increased intensity. Furthermore, fluorescence spectroscopy experiments with Atto-647 conjugated miR-7a (Atto647-miR-7a) and increasing concentrations of the complementary *Klf4* mRNA fragment demonstrated a significant and dose-dependent reduction in fluorescence (Fig. 2F and S 7A). These in vitro studies confirmed the binding of miR-7a to the 3’ UTR region of *Klf4* mRNA, thereby corroborating its direct targeting mechanism.

**Figure 2.**
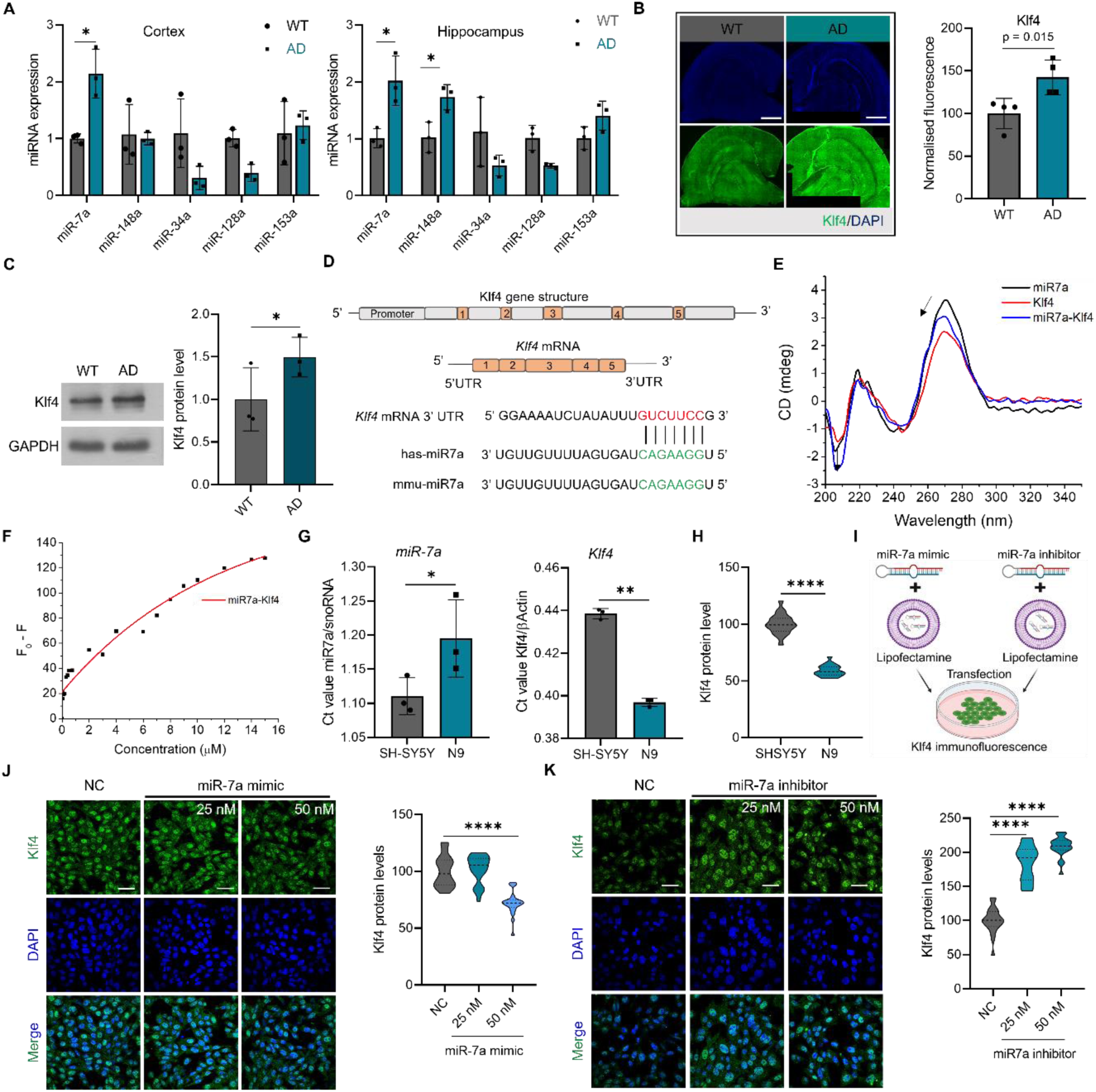
MiR-7a directly regulates increased Klf4 in AD. (**A)** Expression profile of dysregulated miRNAs identified by transcriptomic analysis in cortex and hippocampus of AD mouse brain validated by qRT-PCR (n = 3 biologically independent animals). (**B**) Representative immunofluorescence images of Klf4 in WT and APP/PSEN1 Tg AD mouse brain and its quantification (n = 4 replicates with 2 to 3 section per animal. Scale bar 2 mm). (**C**) Representative western blot images of Klf4 in WT and APP/PSEN1 Tg AD mouse brain and its quantification (n = 3). (**D**) Schematic representation of Klf4 gene and its mRNA with complementary region of miR-7a. (**E**) Circular dichroism spectra of miR-7a, Klf4 and miR-7a-Klf4 annealed in Tri HCl buffer (10 mM, pH 8) performed at 25 °C. (**F**) Dose response curve of change in fluorescence of Atto-647 labelled miR-7a with increasing concentration of *Klf4* mRNA complementary sequence. (**G**) Relative expression of miR-7a and *Klf4* mRNA in SH-SY5Y neuronal and N9 microglial cells by q-RT-PCR and represented Ct_Target_ /Ct_Control_ (n = 3). (**H**) Quantification of Klf4 in SH-SY5Y and N9 cells by immunofluorescence (n > 500 cells quantified for each group. Scale bar 200 μm). (**I**) Schematic representation of transfection experiment. Representative confocal images Klf4 immunofluorescence in SH-SY5Y transfected with miR-7a mimic (**J**) and inhibitor (**K**) with negative control (NC) and its quantification (n > 500 cells quantified for each group). Data are represented as mean ± s.e.m. (**A-C**), (**G**) and (**H**) Unpaired, two-tailed t-test *p < 0.05). (**J**) and (**K**) One-way ANOVA with Bonferroni post hoc test *p < 0.05. Scale bar 200 μm).

We analyzed the expression profiles of miR-7a and *Klf4* through RT-PCR and immunofluorescence in SH-SY5Y neuronal and N9 microglial cell models. The results indicated higher relative expression of miR-7a in N9 microglial cells. In contrast, *Klf4* expression was less pronounced in N9 microglial cells compared to SH-SY5Y cells (Fig. 2G). Subsequent immunofluorescence analysis showed an higher expression of Klf4 protein in SH-SY5Y cells compared to N9, suggesting potential regulation by miR-7a (Fig. 2H and S7B). Klf4, known for its transcription regulation function, is predominantly located in the nucleus. To confirm the direct regulatory effect of miR-7a on Klf4, SH-SY5Y cells were transfected with a miR-7a mimic and inhibitor, along with a negative control, and subjected to immunofluorescence to assess Klf4 levels (Fig. 2I). Results showed a notable decrease in Klf4 expression in cells transfected with the miR-7a mimic, whereas cells with the miR-7a inhibitor exhibited a significant increase in Klf4 protein levels (Fig. 2K). These findings affirm the direct regulatory role of miR-7a on Klf4, emphasizing its potential as an intrinsic ncRNA regulator of Klf4.

### MiR-7a-Klf4 axis regulates AβO mediated microglial activation and neuroinflammation

Alterations in the miR-7a-Klf4 axis observed under AD conditions prompted an investigation into the role of Klf4 in AD-related pathologies. The miRNA-mRNA-Pathway network revealed miRNAs regulatory effect on immune cell pathways within the AD mouse brain. Reports have implicated the role of Klf4 in inflammatory pathways, and AβO are known to trigger microglial activation and neuroinflammation. We examined potential involvement of Klf4 in these processes by monitoring its levels following AβO treatment. N9 microglial cells exhibited dose-dependent increases in Klf4 expression at both mRNA and protein levels upon AβO and lipopolysaccharide (LPS) treatment (Fig. 3A to C, and S8A). Klf4, as a transcription factor, likely governs the expression of proteins that influence microglial state and proinflammatory mediators. To confirm the role of Klf4 in microglial activation and its modulation, we utilized immunofluorescence with Iba1, an activated microglia marker. Post-transfection with a miR-7a mimic and subsequent priming with LPS (500 ng/ml) and AβO (5 μM) challenge, we observed a reduction in microglial activation, indicated by fewer Iba1 positive cells and altered morphology (Fig. 3D and S8B). The activated microglial cells secrete many cytokines that feedback signal microglial cells induce oxidative stress. Specifically, AβO exposure significantly increased intracellular ROS levels, which were mitigated by miR-7a treatment, suggesting a protective effect against ROS production (Fig. 3E).

**Figure 3.**
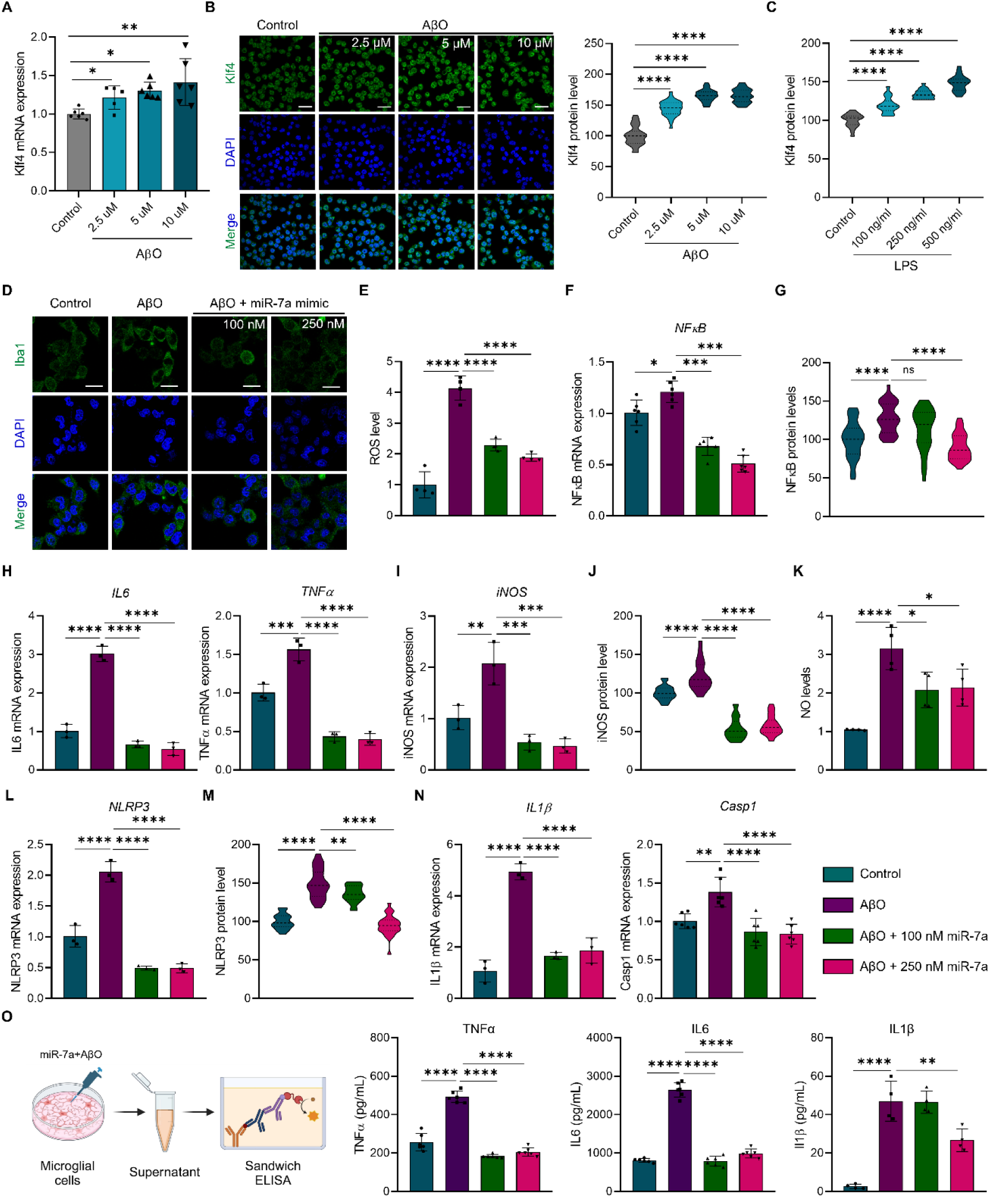
MiR-7a-Klf4 axis regulates microglial activation and neuroinflammation. (**A**) Relative *Klf4* mRNA expression by qRT-PCR of N9 cells treated with increasing concentration of AβO (2.5, 5 and 10 μM) (n = 5 replicates). (**B**) Representative confocal images of Klf4 immunofluorescence in N9 cells treated with increasing concentration of AβO (2.5, 5 and 10 μM) and its quantification (n > 500 cells quantified for each group. Scale bar 200 μm). (**C**) Quantification of Klf4 protein expression in N9 cells treated with increasing concentration of LPS (100, 250 and 500 ng/mL) by immunofluorescence (n > 500 cells quantified for each group. Scale bar 200 μm). (**D**) Representative confocal images of Iba1 immunofluorescence for N9 cells transfected with miR-7a mimic and then challenged with AβO (Scale bar 200 μm). (**E**) Relative intracellular ROS levels detected by DCFDA fluorescence in N9 cells treated with miR-7a mimic and AβO (The bar denotes the normalized mean DCFDA fluorescence detected by well scan with > 80 data points. n = 4 biological replicates.). (**F**) qRT-PCR for *NF-κB* expression of N9 cells treated with miR-7a and AβO (n = 6). (**G**) Quantification of NF-κB protein expression by immunofluorescence (n > 500 cells quantified for each group). RT-PCR of *IL6* and *TNFα* expression (**H**) and *iNOS* (**I**) of N9 cells treated with miR-7a and AβO (n = 3). (**J**) Quantification of iNOS protein expression by immunofluorescence (n > 500 cells quantified for each group). (**K**) Relative NO levels released to media of N9 cells treated with miR-7a and challenged by AβO, quantified by Griess assay (The bar denotes normalized mean absorbance at 540 nm. n = 4 replicates). (**L**) RT-PCR of *NLRP3* expression of LPS primed N9 cells treated with miR-7a and challenged by AβO (n = 3 replicates). (**M**) Quantification of NLRP3 protein expression by immunofluorescence (n > 500 cells quantified for each group). (**N**) RT-qPCR of *IL1β* and *Casp1* expression of LPS primed N9 cells treated with miR-7a and challenged AβO (n =3 and 6 replicates). **o**, Schematic of sandwich ELISA and quantification of TNFα, IL6 and IL1β released in the media of N9 cells treated with miR-7a and challenged with AβO (n= 6 biological replicates. Created with BioRender.com). Data are represented as mean ± s.e.m. All the statistics were performed using One-way ANOVA with Bonferroni post hoc test *p < 0.05).

AβO is known to induce the expression of inflammatory mediators through NF-κB pathway. Further, we explored the regulatory impact of miR-7a-Klf4 axis on the NF-κB levels (Fig. 3F, G and S8C). MiR-7a-mediated Klf4 silencing resulted in decreased *NF-κB* expression, corroborated by the downregulation of proinflammatory cytokines *IL6* and *TNFα* (Fig. 3H). We assessed the expression levels of iNOS and found that miR-7a treatment to target Klf4 has effectively reduced iNOS mRNA and protein expression in AβO treated microglial cells (Fig. 3I, J and S8D). Further determination of relative NO by Griess assay has demonstrated the significant reduction of NO levels (Fig. 3K). We also assessed NLRP3 expression, a key player in neuroinflammatory pathways, and found it significantly elevated following AβO treatment (Fig. 3L, M and S8E). However, miR-7a treatment substantially decreased NLRP3 expression and its downstream cytokines IL1β and caspase 1 (Fig. 3N). The mitigation of neuroinflammation was further confirmed by quantifying neuroinflammatory mediators TNFα, IL6, and IL1β via sandwich ELISA. The results demonstrated that miR-7a treatment targeting Klf4 effectively inhibited the secretion of proinflammatory mediators induced by AβO (Fig. 3O). Overall, our studies indicate that miR-7a mimic treatment targeting Klf4 significantly reduces microglial activation and attenuates various neuroinflammatory pathways, including NF-κB, iNOS, and NLRP3 inflammasome, thereby alleviating neuroinflammation. The disruption of Klf4-mediated neuroinflammation by miR-7a suggests its potential as an RNA therapeutic targeting neuroinflammation in AD.

### MiR-7a-Klf4 axis regulates neuronal ferroptosis

Ferroptosis, characterized by iron-mediated lipid peroxidation and cell death, is implicated in neurodegenerative disease pathogenesis. Klf4, a mediator of apoptotic cell death and regulator of iron homeostasis, led us to hypothesize its role in ferroptosis. We investigated this hypothesis using RAS-selective lethal 3 (RSL3)-induced ferroptosis in SH-SY5Y neuroblastoma cells, to understand the role of miR-7a-Klf4 axis. RSL3 inhibits glutathione peroxidase 4 (GPX4), essential for ferroptosis, causing neuronal cell death. Upon ferroptosis induction by RSL3, Klf4 mRNA and protein levels significantly increased, suggesting the regulatory role of Klf4 in ferroptosis (Fig. 4A and B). As Klf4 being transcription factor involved in various biological processes, its upregulation potentially mediate ferroptotic cell death. If Klf4 influences ferroptosis, its suppression by miR-7a mimic treatment could protect cells from ferroptotic cell death. To confirm this, we assessed different hallmarks of ferroptosis in RSL3-induced conditions and treatment with a miR-7a mimic. Neuronal cells were transfected with a miR-7a mimic, then challenged with RSL3 and analyzed for ferroptosis hallmarks to observe the rescue effect.

**Figure 4.**
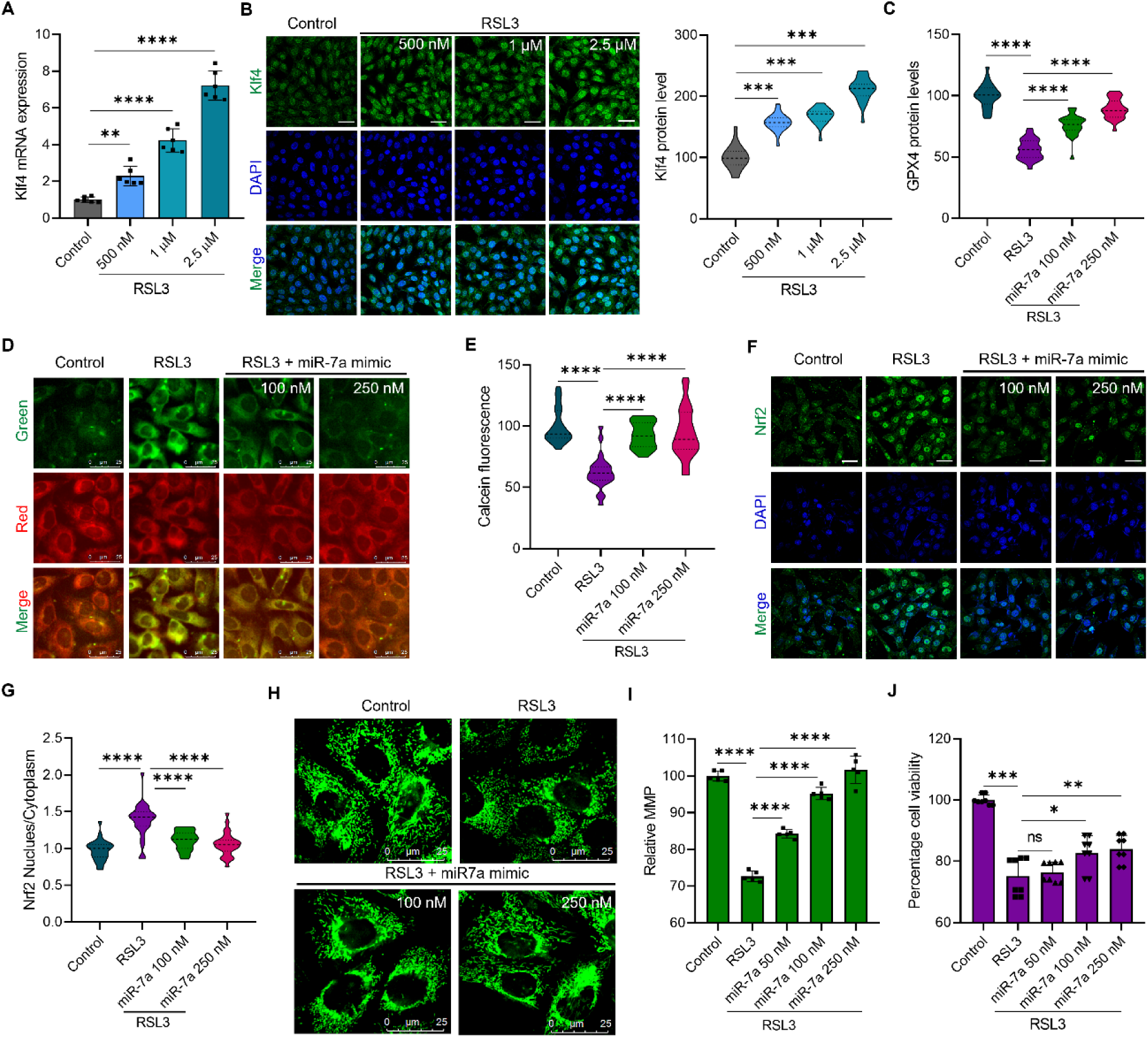
MiR-7a-Klf4 axis regulates neuronal ferroptosis. (**A**) Relative *Klf4* mRNA expression by qRT-PCR of SH-SY5Y cells treated with increasing concentration of RSL3 (0.5, 1 and 2.5 μM) (n = 6 replicates). (**B**) Representative confocal images Klf4 immunofluorescence in SH-SY5Y cells treated with increasing concentration of RSL3 (0.5, 1 and 2.5 μM) and its quantification (n > 500 cells quantified for each group. Scale bar 200 μm). (**C**) Quantification of GPX4 protein expression by immunofluorescence (n > 500 cells quantified for each group). (**D**) Fluorescence images of C11-Bodipy lipid peroxidation probe (Scale bar 25 μm). (**E**) Relative Fe^+3^ levels quantified by Calcein-AM fluorescence imaging (n > 500 cells quantified for each group). Representative confocal images of Nrf2 immunofluorescence (**F**) and quantification of its translocation to nucleus (**G**) (The bar denotes mean of nuclear/cytoplasmic fluorescence. n > 500 cells quantified for each group). (**H**) Mitochondrial imaging using Mitotracker green (Scale bar 25 μm). (**I**) Relative MMP quantified using Rho123 potential dependent fluorescent dye (Normalized mean Rho123 fluorescence detected by well scan with > 80 data points. n = 5 replicates). (**J**) Cell viability quantified by MTT assay (n = 8 biological replicates). Data are represented as mean ± s.e.m. All the statistics were performed using One-way ANOVA with Bonferroni post hoc test *p < 0.05).

We assessed GPX4 levels, a marker known to reduce during RSL3-induced ferroptotic cell death, via immunofluorescence assay. Results revealed a significant reduction in GPX4 levels following RSL3 treatment, which was dose-dependently rescued in cells treated with a miR-7a mimic (Fig. 4C and S9A). The lipid peroxidation, a ferroptosis hallmark, was monitored using a red fluorescent C11-Bodipy probe, which emits green fluorescence upon oxidation by lipid peroxides. The Bodipy dye fluorescence imaging showed increased lipid peroxide levels in RSL3-treated cells, while miR-7a treatment prevents lipid peroxidation (Fig. 4D and S9B). Quantitative analysis in well plate assay confirmed a significant rise in lipid peroxidation, reduced significantly by miR-7a treatment. Ferroptosis is initiated by an intracellular labile iron pool (LIP) through Fenton-type reactions causing lipid peroxidation. Employing calcein-AM fluorescence imaging, we analyzed LIP under ferroptosis condition and its mitigation by miR-7a mimics. A marked reduction in calcein-AM fluorescence indicated increased LIP in RSL3-treated cells, whereas miR-7a treated cells retainedfluorescence, implying inhibition of iron overload (Fig. 4E and S9C). Iron-triggered Fenton-type reactions generate ROS, leading to oxidative stress and Nrf2 activation. Measuring intracellular ROS using 2’-7’-dichlorodihydrofluorescein diacetate (DCFDA), we found miR-7a significantly reduced ROS in RSL3-challenged neuronal cells (Fig. S9D). Under oxidative stress, Nrf2 translocate to the nucleus, inducing antioxidant gene expression. The reduction of LIP and ROS, coupled with GPX4 rescue by miR-7a, potentially prevents oxidative stress and Nrf2 activation (Fig. 4F, 4G and S9E). Excessive oxidative stress and lipid peroxidation can cause mitochondrial damage in ferroptosis. We evaluated the effect of miR-7a-mediated Klf4 silencing in preventing mitochondrial structural and functional damage. Mitotracker green imaging revealed mitochondrial fragmentation, as observed by the fragmented and bulged mitochondria in ferroptotic cells, while miR-7a-treated cells retained normal long tubular mitochondrial morphology, indicating prevention of mitochondrial damage (Fig. 4H). To ascertain functional impairment, we quantified the relative mitochondrial membrane potential (MMP) utilizing rhodamine 123 (Rho123), a fluorescent probe whose signal intensity is dependent on MMP. In the context of RSL3-induced ferroptosis, the membrane potential reduced to 75% compared to the control. Contrastingly, miR-7a-treated cells exhibited a restoration of membrane potential in a dose-dependent manner, achieving full recovery at a concentration of 250 nM (Fig. 4I).

The attenuation of Klf4 expression via miR-7a mimic intervention was instrumental in reversing the hallmarks of ferroptosis, underscoring pivotal role of Klf4 in neuronal ferroptosis. Encouraged by the protective effect of miR-7a, we proceeded to examine its efficacy in safeguarding neuronal cells from ferroptotic death. The cell viability assays showed a significant amelioration of ferroptotic cell death attributed to miR-7a-facilitated Klf4 suppression (Fig. 4J). These findings highlight the significant role of Klf4 in ferroptosis and demonstrate the therapeutic promise of miR-7a as an RNA-based agent targeting ferroptosis in AD and other neurodegenerative conditions.

### Pharmacological targeting of miR-7a-Klf4 axis using small molecules to rescue neuroinflammation and ferroptosis

The delivery of RNA therapeutics to the brain for treating CNS disorders is a formidable challenge, emphasizing the necessity for crafting small molecule modulators that can precisely manipulate the regulatory axis to enhance therapeutic outcomes. The strategic inhibition of Klf4 by miR-7a has demonstrated efficacy in mitigating microglial activation, neuroinflammation, and ferroptotic neuronal death, which are pivotal in the AD pathology. Development of RNA therapeutics or small molecules that can modulate the miR-7a-Klf4 axis either by bolstering endogenous miR-7a levels or by directly targeting Klf4 mRNA and protein represents a promising therapeutic avenue for AD and other neurodegenerative disorders (Fig. 5A).

**Figure 5.**
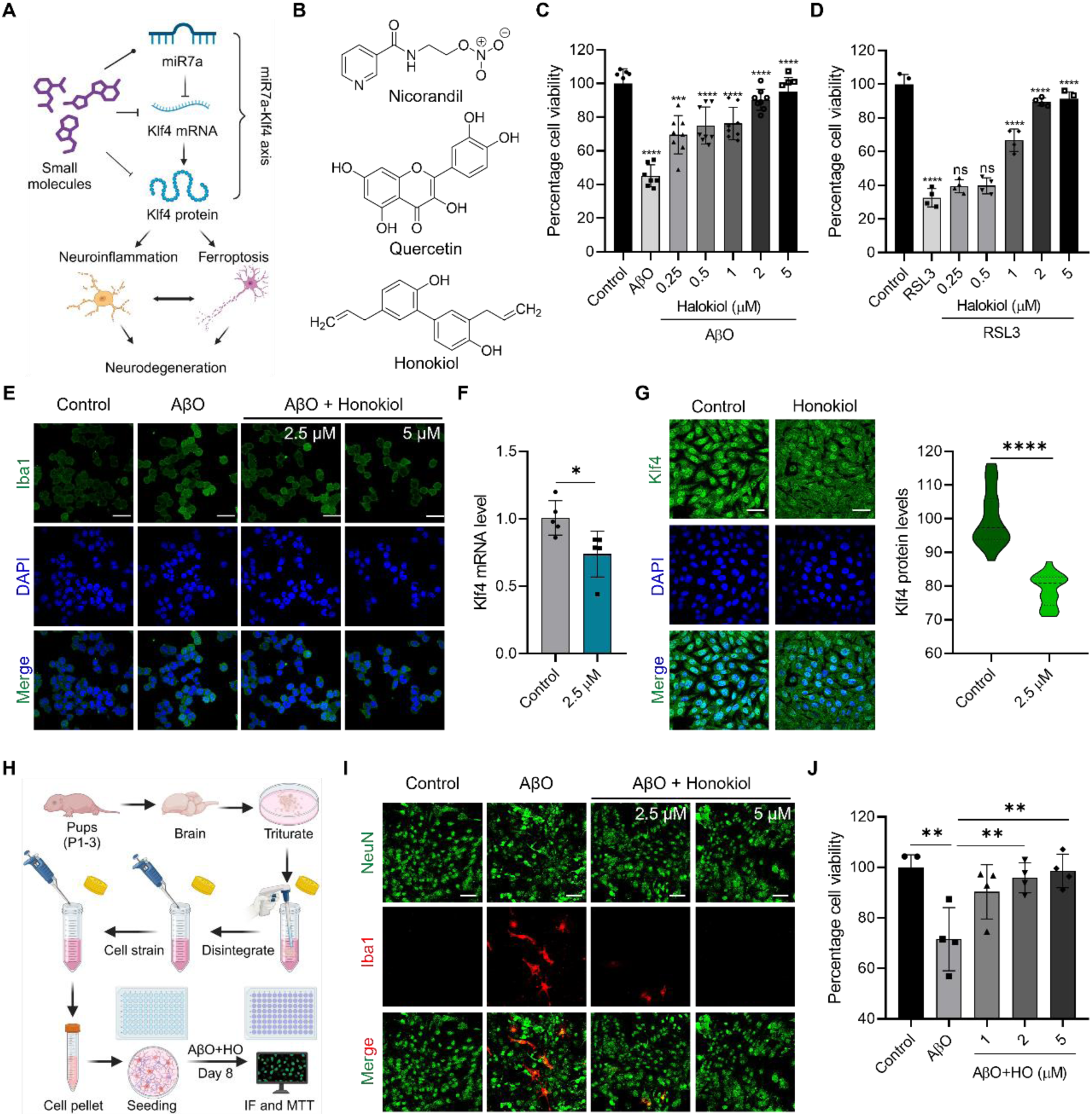
Pharmacological targeting of miR-7a-Klf4 axis rescue neuroinflammation and ferroptosis. (**A**) Schematic representation of chemical targeting of miR-7a-Klf4 axis to curb neuroinflammation and ferroptosis. (**B**) Chemical structures of small molecules used to modulate miR-7a-Klf4 axis. (**C**) N9 microglial cell rescue from AβO induced inflammatory cell death by honokiol (HO) in dose dependent manner (n = 8). (**D**) SH-SY5Y neuronal cell rescue from RSL3 induced ferroptotic cell death by honokiol in dose dependent manner (n = 8). (**E**) Representative confocal images of Iba1 immunofluorescence of N9 cells treated with AβO and honokiol (Scale bar 200 μm). (**F**) Relative *Klf4* mRNA expression by qRT-PCR of SH-SY5Y cells treated with honokiol (2.5 μM) (n = 5. Unpaired, two-tailed t-test p < 0.05). (**G**) Representative confocal images of Klf4 immunofluorescence in SH-SY5Y cells treated with honokiol (2.5 μM) and its quantification (n > 500 cells quantified for each group. Scale bar 200 μm). (**H**) Schematic showing the protocol for mouse primary neuron-glia mixed culture (Created with BioRender.com). (**I**) Representative confocal images of primary neuron-glia mixed culture treated with AβO (2.5 μM) alone and in presence of honokiol (2.5 and 5 μM) and probed for NeuN and Iba1 markers for neuronal cells and activated microglial cells (Scale bar 200 μm). (**J**) Cell viability of primary neuron-glia mixed culture treated with AβO (2.5 μM) and honokiol (2.5 and 5 μM) measured by MTT assay (n = 4). Data are represented as mean ± s.e.m. Statistics used for (**F**) and (**G**) is unpaired, two-tailed t-test p < 0.05. For (**C**), (**D**) and (**J**), One-way ANOVA with Bonferroni post hoc test *p < 0.05.

**Figure 6.**
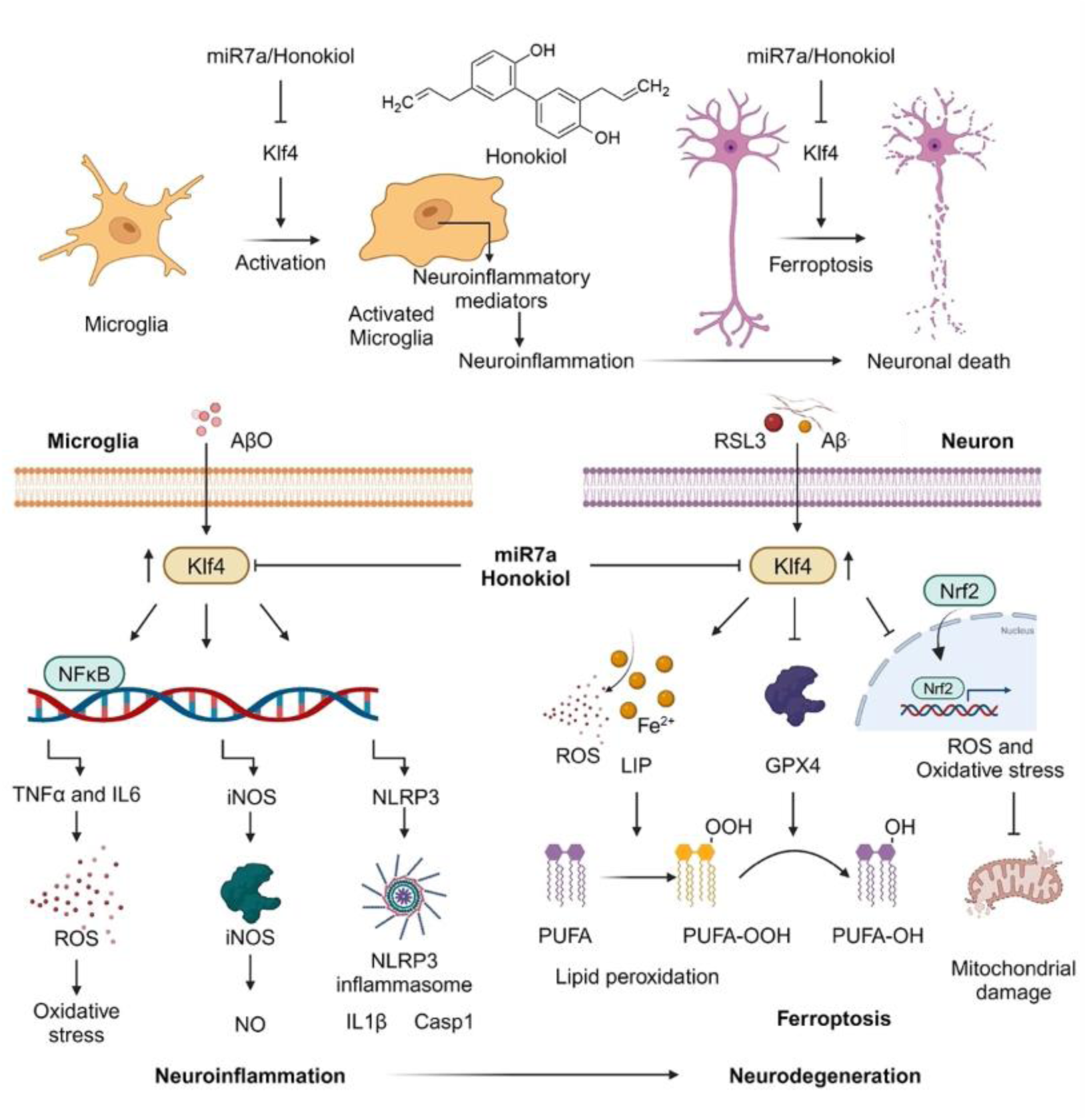
Schematic representation of miR-7a-Klf4 axis regulating neuroinflammation and ferroptosis resulting in neurodegeneration through various pathways and its targeting by using miR7a and honokiol (Created with BioRender.com).

A few small molecules have been reported to elevate miR-7a levels and concurrently suppress Klf4 expression. We evaluated such molecules namely nicorandil, quercetin, and honokiol, potentially modulate the miR-7a-Klf4 axis (Fig. 5B). We assessed their efficacy in mitigating neuroinflammatory and ferroptotic cell death via cell viability assays. Notably, honokiol demonstrated a concentration-dependent effect in rescuing cells from neuroinflammation and ferroptosis compared to others (Fig. 5C, D and S10A to D). The BBB permeability of honokiol and its minimal cytotoxicity at concentrations up to 20 μM in SH-SY5Y cells and 10 μM in N9 cells underscore its therapeutic potential (Fig. S10E and F). Honokiol was effective at lower concentrations in alleviating neuroinflammation and ferroptosis. Further honokiol effectively rescued N9 microglial cells from AβO mediated activation as evident from Iba1 immunofluorescence (Fig. 5E). The expression analysis of Klf4 mRNA and protein revealed that honokiol significantly reduced the Klf4 expression at both levels (Fig. 5F and G). Targeting Klf4 rescued neuroinflammation and ferroptosis, that suggest Klf4 as a potential therapeutic target for AD and other neurodegenerative disorders.

To confirm the rescue effect of honokiol on neuroinflammation and neuronal death, we have employed mice primary neuron-glia mixed culture model that mimics in vivo conditions. We have isolated cells from day 1 pup (P1) brain and cultured for 8 days followed by treatments (Fig. 5H). Primary mixed cell culture was primed with LPS (250 ng/mL) for 4 h and then challenged with AβO (2.5 μM) for 24 h. Cells were subjected to NeuN and Iba1 immunostaining to probe neuron and activated microglial cells, respectively. The confocal images show the activation of microglia in AβO treatment, which was decreased by honokiol treatment (Fig. 5I). The activated microglia have resulted in decreased neuronal population indicating the neuronal death that was rescued by honokiol. We have performed cell viability assay to confirm the rescue of AβO induced primary neuronal cell death by honokiol. The results show concentration dependent rescue of cell death suggesting its therapeutic effect (Fig. 5J). Further research is necessary to explore the potential of honokiol and its derivatives to effectively modulate AD and other neurodegenerative conditions.

## DISCUSSION

AD is a complex neurodegenerative disorder and the key regulatory mechanisms driving its onset and progression are not fully understood. As upstream transcription regulators, miRNA target multiple mRNAs across cellular pathways, orchestrating cellular homeostasis and disease pathways. The multitargeting capability of miRNAs establishes a miRNA-mRNA regulatory network, enabling the precise regulation of cellular pathways. Despite the critical regulatory role of miRNAs, their involvement in AD and the potential miRNA-mRNA regulatory networks remain underexplored. The failure to develop effective therapeutics for AD underscores the need to decipher the miRNA-mRNA-pathway network implicated in AD pathology, which could reveal new diagnostic and therapeutic targets. In this study, we utilized an APP/PSEN1 Tg AD mouse model to profile miRNA expression alterations in the AD brain, identifying significant changes in the cortex and hippocampus. Independent clustering of miRNA profiles from WT and AD mouse brains highlighted a substantial dysregulation, with 21 and 11 miRNAs significantly altered in the cortex and hippocampus, respectively. This transcriptomic landscape revealed both upregulated and downregulated miRNAs, reprogramming the miRNA-mRNA pathway network and contributing to AD pathology. By constructing a miRNA-mRNA-pathway and disease network based on validated mRNA target predictions, we observed that miRNAs regulate various altered disease pathways in AD. Modulating the miRNA-mRNA axis may offer a strategy to reprogram this network, leading to improved disease-modifying treatments. Gene ontology and functional over-representation analyses identified pathways related to stress response, immune function, cell death and cycle, neuronal and synaptic regulation, protein homeostasis, and other processes linked to CNS functions, innate immunity, and neurodegeneration. Further examination of the miRNA-mRNA-disease network revealed the influence of altered miRNAs in various CNS and neurodegenerative disorders, including AD. Our analysis underscores the significance of the miRNA regulatory network in disease pathology and neuroprotective role of miRNAs in AD and other neurodegenerative diseases.

The transcriptomic analysis, corroborated by qPCR experiments, indicates an upregulation of miR-7a in the cortex and hippocampus of AD mouse brain. MiR-7a, an abundantly expressed miRNA in the brain and evolutionarily conserved, is recognized for its neuroprotective properties and influence on CNS functions [38]. It targets mRNAs associated with AD and Parkinson disease (PD), such as IDE and a-Syn [39, 40]. Reports suggest the regulatory effect of miR-7a on Klf4, a pleotropic transcription factor implicated in cancer and AD. Elevated Klf4 expression in AD brains suggests its potential involvement in disease onset and progression. We propose that elevated Klf4 in AD brains serves as a principal transcriptional regulator of AD-related pathological pathways and miR-7a targeting it for neuroprotection. MiR-7a expression aims to counterbalance overexpression of Klf4 in AD, establishing a miR-7a-Klf4 axis that play role in neurodegeneration. We have shown the direct interaction of miR-7a with Klf4 at the 3’ UTR region via CD and fluorescence spectroscopy. Klf4 mRNA contains two miR-7a binding sites, one conserved among mammals and another across vertebrates [38]. The inverse expression patterns of miR-7a and *Klf4* mRNA in SH-SY5Y neuroblastoma and N9 microglial cells imply the direct regulatory role of miR-7a. Transfection assays with miR-7a mimic and inhibitor affirm the regulation of Klf4 by miR-7a and the significance of miR-7a-Klf4 axis in AD pathology. The role of the miR- 7a-Klf4 axis has been studied in cancer cells and its role in regulating various pathways linked to cancer development, progression and metastasis, and resistance [41–44]. The direct regulation of Klf4 by miR-7a and its role in controlling various pathways and cancer pathobiology suggest role of miR-7a-Klf4 axis in neurons and microglial pathology linked to AD and other neurodegenerative disorders.

The miRNA-mRNA-pathway analysis has revealed alterations in immune cell pathways within the brain affected by AD. Notably, Klf4 has been implicated in microglial activation and neuroinflammation. Our study demonstrated that Klf4 expression is induced at both mRNA and protein levels in LPS and AβO treated microglial cells. The activation states of microglia and various neuroinflammatory pathways are ameliorated when cells are transfected with miR-7a mimic and challenged with AβO. These findings underscore the pivotal role of the miR-7a-Klf4 axis in microglial activation and neuroinflammation, particularly in regulating NF-κB, iNOS, and NLRP3 pathways. There are reports suggesting the induction of Klf4 expression in LPS and AβO treated microglial cells [45–47]. Elevated Klf4 expression has been observed in activated microglial cells, both in vitro and in vivo, using the 5xFAD mouse model [48]. However, there is a dearth of information regarding the intrinsic regulator of Klf4 under neuroinflammatory conditions and the specific downstream pathways it regulates. Our observations indicate that Klf4 targets NF-κB expression and downstream proinflammatory mediators such as TNFα and IL6. Similar observations were reported earlier wherein Klf4 induces NF-κB signaling [49] and Klf4 directly interacts with NF-κB and regulates its activity [50]. Contradictory reports exist, with some suggesting that Klf4 is downregulated in activated glial cells exerts an anti-inflammatory effect by suppressing the NF-κB pathway in macrophages, endothelial, microglial cells and astrocytes [50–54]. The inhibition of MEG3, a lncRNA mediated suppression of Klf4 induced inflammation in microglial cells of cerebral ischemia-reperfusion injury model [52]. It is noteworthy that Klf4, as a pleiotropic transcription factor, can regulate NF-κB signaling both positively and negatively, depending on the cell type and disease context. Klf4 directly regulates *iNOS* expression, leading to an increase in NO production [45]. Suppression of Klf4 reduces NLRP3 expression and downstream mediators such as IL1β and caspase 1 [55]. Additionally, NF-κB is also known to regulate NLRP3 expression and Klf4 regulates both NF-κB and NLRP3 that accounts for significant reduction of NLRP3 and downstream IL1β and Caspase 1 levels. NLRP3 is a direct target for miR-7a and inhibited microglial NLRP3 inflammasome activation in PD model [56]. Further, Klf4 binds to the NLRP3 promoter and positively regulates its expression. Other miRNAs, including miR-146a-5p and miR-92a, have been reported to target Klf4, suggesting the regulatory role of the miR-Klf4 axis in neuroinflammation [57, 58].

Klf4 is recognized for its key role in cell death mechanisms, including apoptosis, oxidative stress, and iron homeostasis [59]. The miR-mRNA-pathway analysis indicates significant alterations in cellular stress responses, as well as cell cycle and death pathways. We have observed a marked induction of Klf4 in neuronal ferroptosis and upregulation of Klf4 appears to orchestrate several processes involved in ferroptosis. Inhibition of Klf4 in ferroptotic cells using miR-7a mimics leads to a reduction in ferroptotic markers such as Fe^3+^, GPX4, lipid peroxidation, oxidative stress, Nrf2 signaling, and mitochondrial damage, thereby enhancing cell survival. Previous studies have highlighted the role of the miR-7a-Klf4 axis in regulating oxidative stress and apoptosis mediated neuronal cell death [60, 61]. Our findings reveal that miR-7a mimic treatment reduces ROS and oxidative stress, suggesting the involvement of Klf4 in redox homeostasis. It also prevents the accumulation of Fe^3+^ in neuronal cells challenged with RSL3, possibly through the regulation of heme uptake mediated by heme carrier protein 1 (HCP1) [62]. Moreover, miR-7-5p has been reported to downregulate the expression of iron-related genes such as transferrin receptor (TfR), ferritin, and iron regulatory protein 2 (IRP2), as well as ferroptosis markers like lipid peroxidation (ALOX12) [63]. Our study reveals that miR-7a modulates iron levels and lipid peroxidation in ferroptosis by targeting Klf4, suggesting that the miR-7a-Klf4 axis regulates ferroptosis by regulating LIP, ROS-oxidative stress, mitochondrial integrity, and lipid peroxidation in neuronal cells.

Investigations into the involvement of the miR-7a-Klf4 axis in neuroinflammation and ferroptosis have led us to identification of small molecule modulators to counteract these pathological pathways. Identification of small molecules potentially induce miR-7a is one of the approaches to target miR7a-Klf4 axis [64]. Nicorandil has been shown to significantly increase miR-7a levels, offering relief from astrocytic inflammation [65]. Honokiol, on the other hand, diminishes Klf4 protein levels and NF-κB, curtailing the expression of iNOS, Cox- 2, MCP-1, IL-6, and TNF-α in both LPS-treated microglia and mouse brains [66]. Similarly, quercetin targets Klf4, safeguarding SH-SY5Y cells from H_2_O_2_-induced oxidative stress and apoptosis [60]. We employed these molecules to pharmacologically modulate the miR-7a- Klf4 axis, assessing their effects on neuroinflammatory-driven microglial cell death and RSL3- induced neuronal ferroptosis. Our results indicate that quercetin effectively alleviates neuroinflammation, while honokiol rescues cells from both neuroinflammation and ferroptosis at lower doses due to its ability to reduce Klf4 expression. Notably, higher concentrations of honokiol have been associated with the induction of ferroptosis, a phenomenon warranting further exploration to understand its concentration-dependent dual role [67, 68]. Comprehensive study evaluating the therapeutic potential of honokiol in mouse primary mixed culture have confirmed the role of miR-7a-Klf4 axis in neuroinflammation and neurodegeneration. Identifying and assessing new miRNA and small molecule therapeutics that modulate this axis is crucial for developing improved treatments for AD and other neurodegenerative disorders [64, 69]. These findings highlight the potential for novel therapeutic interventions that target miR-7a-Klf4 axis, enhancing the diseases management strategies.

In summary our study has unveiled a spectrum of miRNAs that exhibit altered expression patterns in AD, leading to the establishment of a reprogrammed miRNA-mRNA-Pathway network pertinent to AD. Notably, we have discovered an elevated level of Klf4 and induction of miR-7a in the AD brain to target Klf4 for neuroprotection. The miR-7a-mediated attenuation of Klf4 elucidates its pivotal role in orchestrating neuroinflammation via diverse molecular conduits, including NF-κB signaling, iNOS, and the NLRP3 inflammasome cascade. Furthermore, our study highlighted the significant role of the miR-7a-Klf4 axis in ferroptosis- a distinct modality of cell death characterized by LIP overload, lipid peroxidation, ROS, oxidative stress, and mitochondrial impairment. The strategic pharmacological manipulation of the miR-7a-Klf4 axis utilizing small molecule has demonstrated efficacy in ameliorating neuroinflammation and ferroptosis, positing the miR-7a-Klf4 axis as a viable pharmacological target for AD and potentially other neurodegenerative disorders. These insights pave the way for the inception of novel therapeutic modalities aimed at mitigating the impact of these chronic diseases.

## DATA AVAILABILITY

NanoString miRNA transcriptomic data is deposited in NCBI GEO repository with accession number GSE271276 and will be made public upon acceptance of manuscript. All data supporting the findings of this study are available within the paper and Supplementary files. All other data generated in this study are available from the corresponding author

## SUPPLEMENTARY DATA

Supplementary Data are available at NAR online.

## AUTHOR CONTRIBUTIONS

Conceptualization: TG and MR, Methodology: MR and TG, Investigation and Visualization: MR, Supervision: TG, Writing - original draft: MR, Writing, review & editing: TG and MR.

## ACKNOWLEDGEMENTS

We thank Theracues Innovations Pvt Ltd, Bangalore, India, a service provider for NanoString platforms, for Bioinformatic analysis. We thank Shivprasad P S for help in confocal imaging, lab members and Prathamesh for discussion. MR thanks UGC and JNCASR for the student fellowships.

## FUNDING

The authors thank JNCASR, DST (CEFPRA grant: IFCPAR/CEFIPRA-62T10-3), core grant (CRG/2020/004594), Science and Engineering Research Board (SERB), New Delhi, India, for the funding.

## CONFLICT OF INTEREST

All other authors declare they have no competing interests.

## Notes

### Competing Interest Statement

The authors have declared no competing interest.

