## Supplementary figures and images for "MiR-7a-Klf4 axis as a regulator and therapeutic target of neuroinflammation and ferroptosis in Alzheimer’s disease"

### Suplementary Data 1_Cortex_miRNA-MRNA-pathway network.pdf

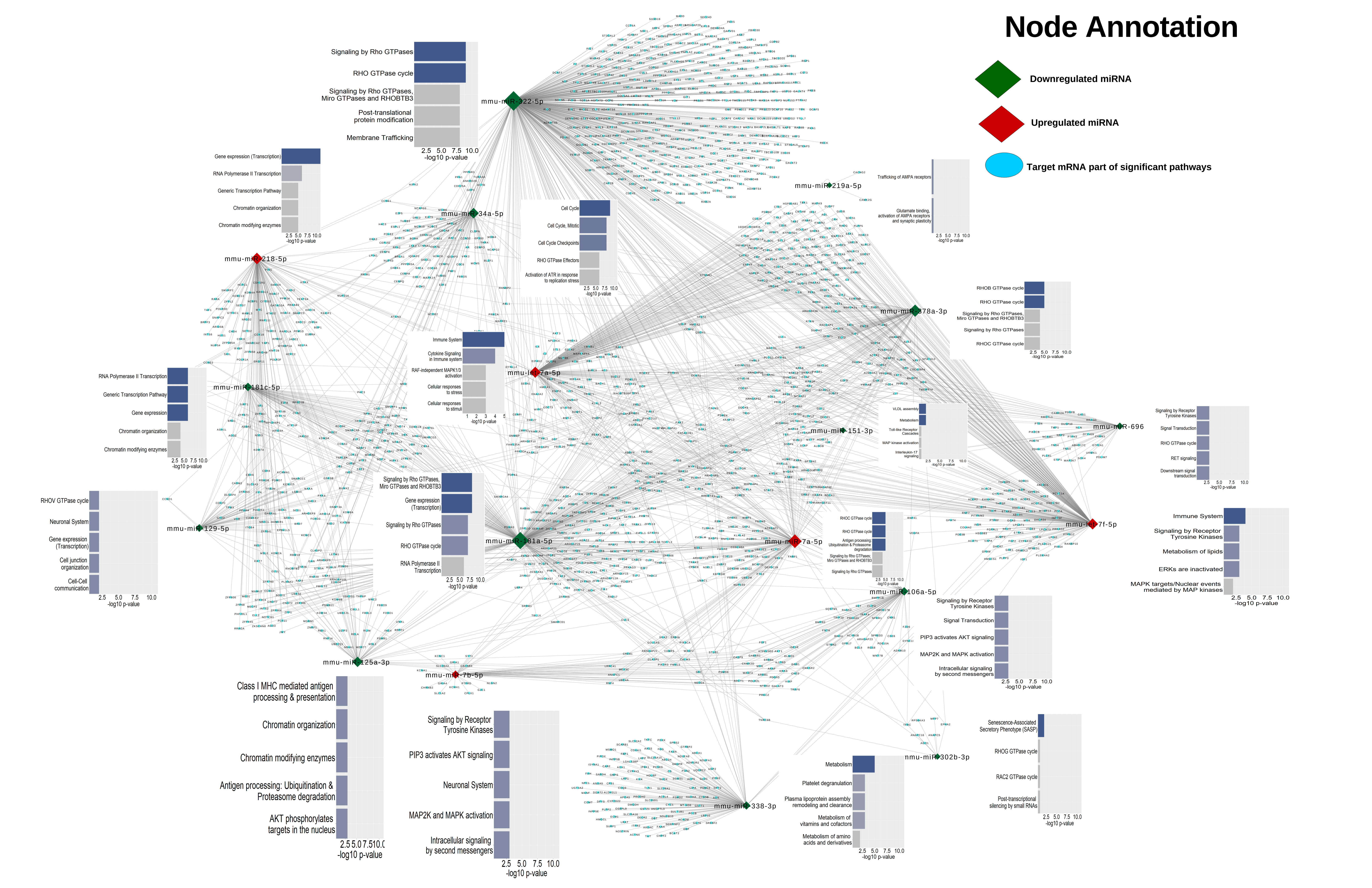

### Suplementary Data 2_Hippocampus_miRNA-MRNA-pathway network.pdf

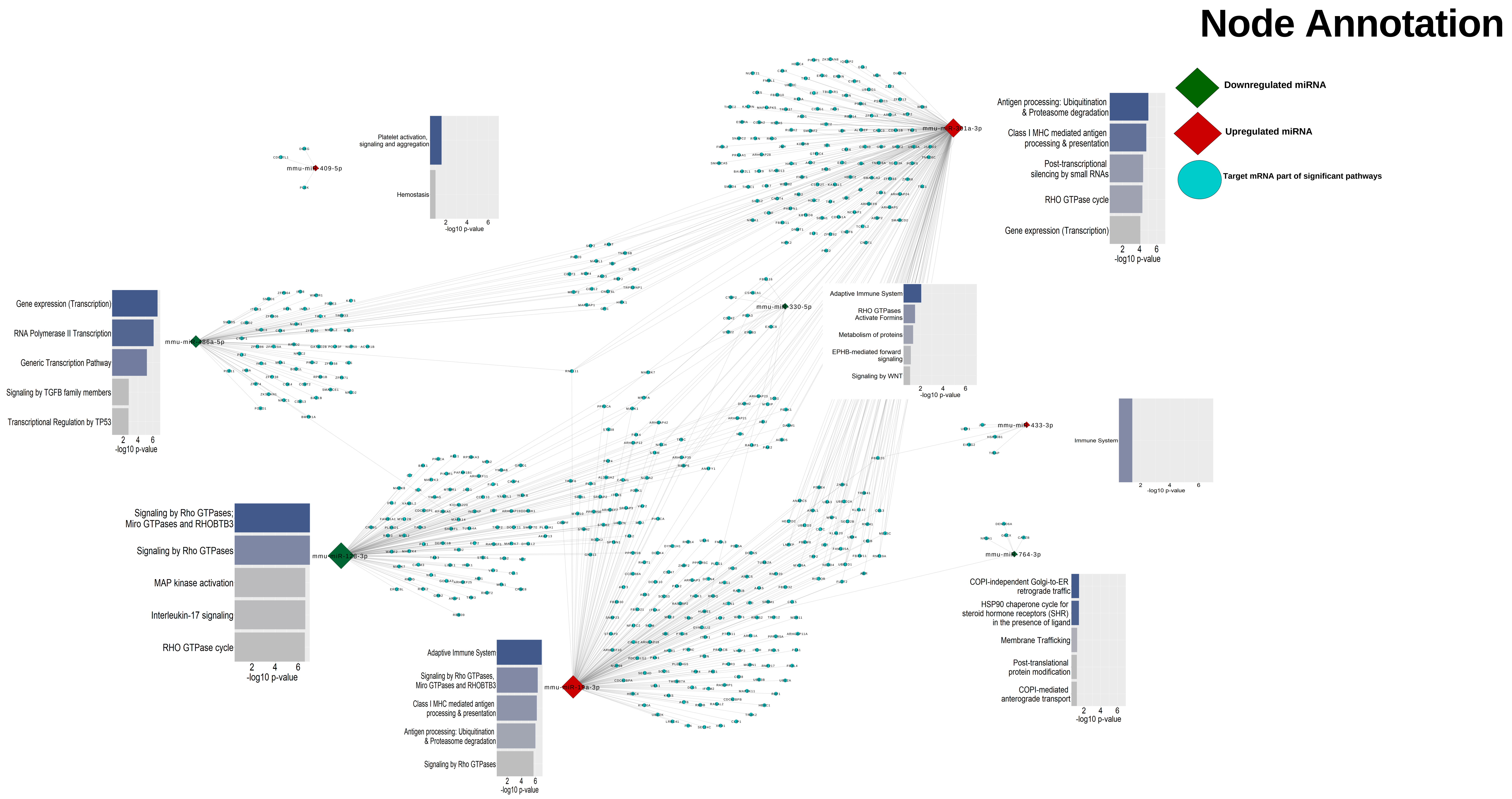

### Suplementary Data 3_Western blots.pdf

Raw western blots of Klf4 and GAPDH

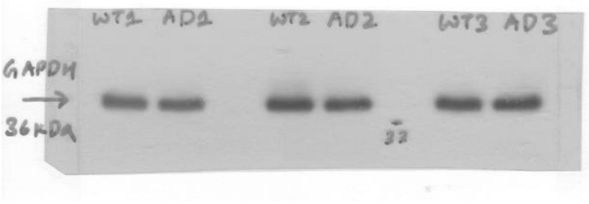


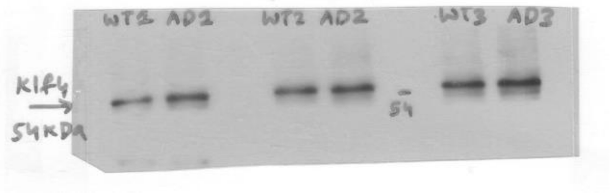
