## Supporting Information for "MiR-7a-Klf4 axis as a regulator and therapeutic target of neuroinflammation and ferroptosis in Alzheimer’s disease"

**Content**

Figs. S1 to S10

Tables S1

**Other Supplementary DAta for this manuscript include the following**

Data S1 to S3

Tables S2 to S10

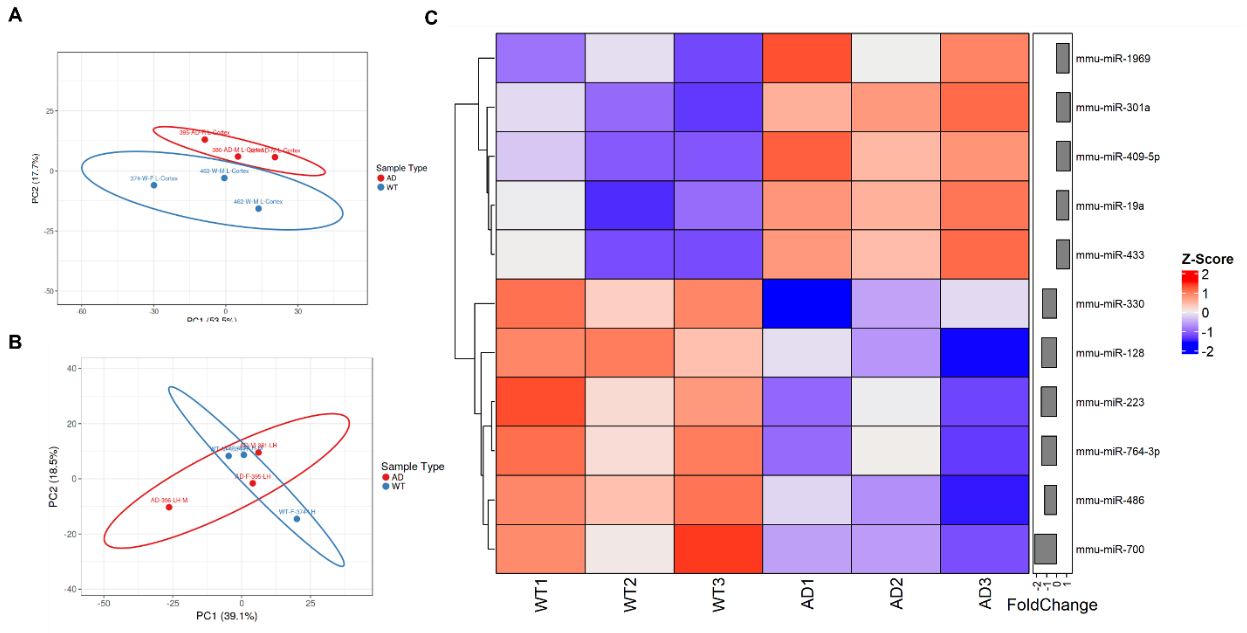


**Figure S1.** Differentially expressed miRNAs and altered miRNA-mRNA-pathway/disease network. The multidimensional scaling plot of cortex (**A**) and hippocampus (**B**) from APP/PSEN1 Tg AD mouse brain miRNA profiling revealed a stronger correlation and clustering of WT and AD samples. (**C**) Heatmap of the top variable miRNAs of hippocampus generated from miRNA transcriptomics.

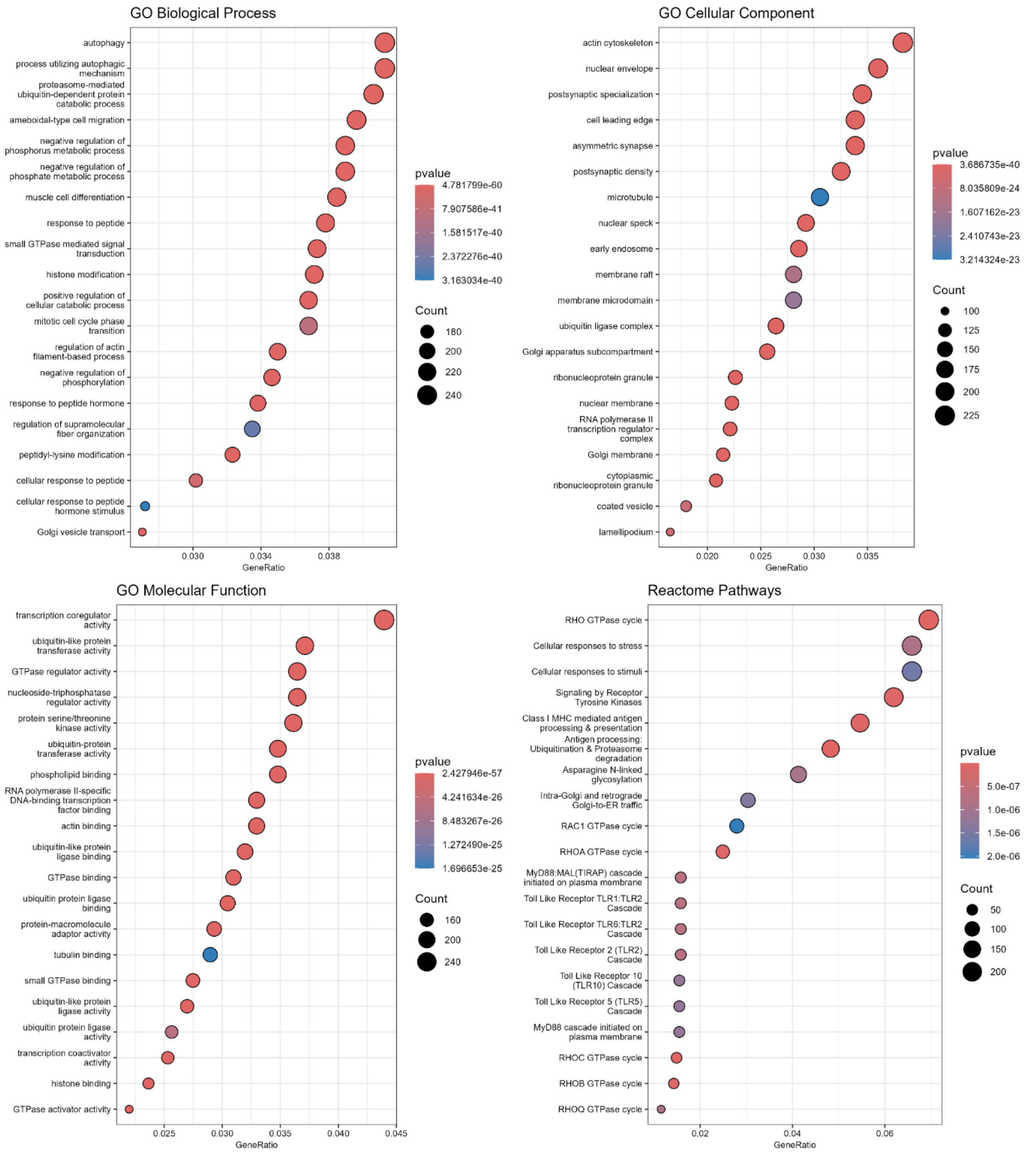


**Figure S2.** Functional overrepresentation analysis for identified mRNA targets of downregulated miRNA in cortex to identify biological processes, cellular components, molecular function and reactome pathways regulated by downregulated miRNAs ($p < 0.05$).

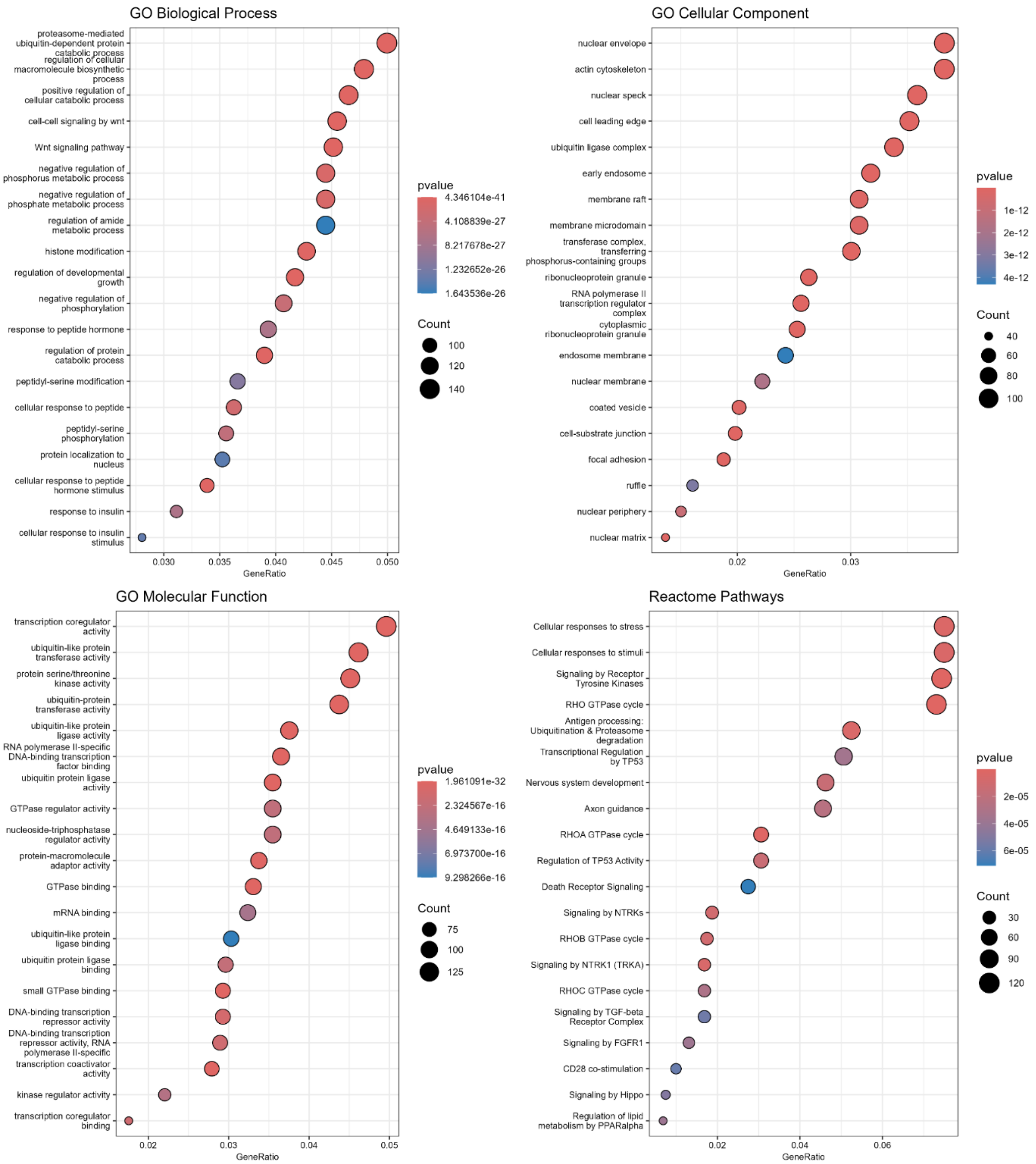


**Figure S3.** Functional overrepresentation analysis for identified mRNA targets of upregulated miRNA in cortex identifying biological processes, cellular components, molecular function and reactome pathways regulated by upregulated miRNAs ($p < 0.05$).

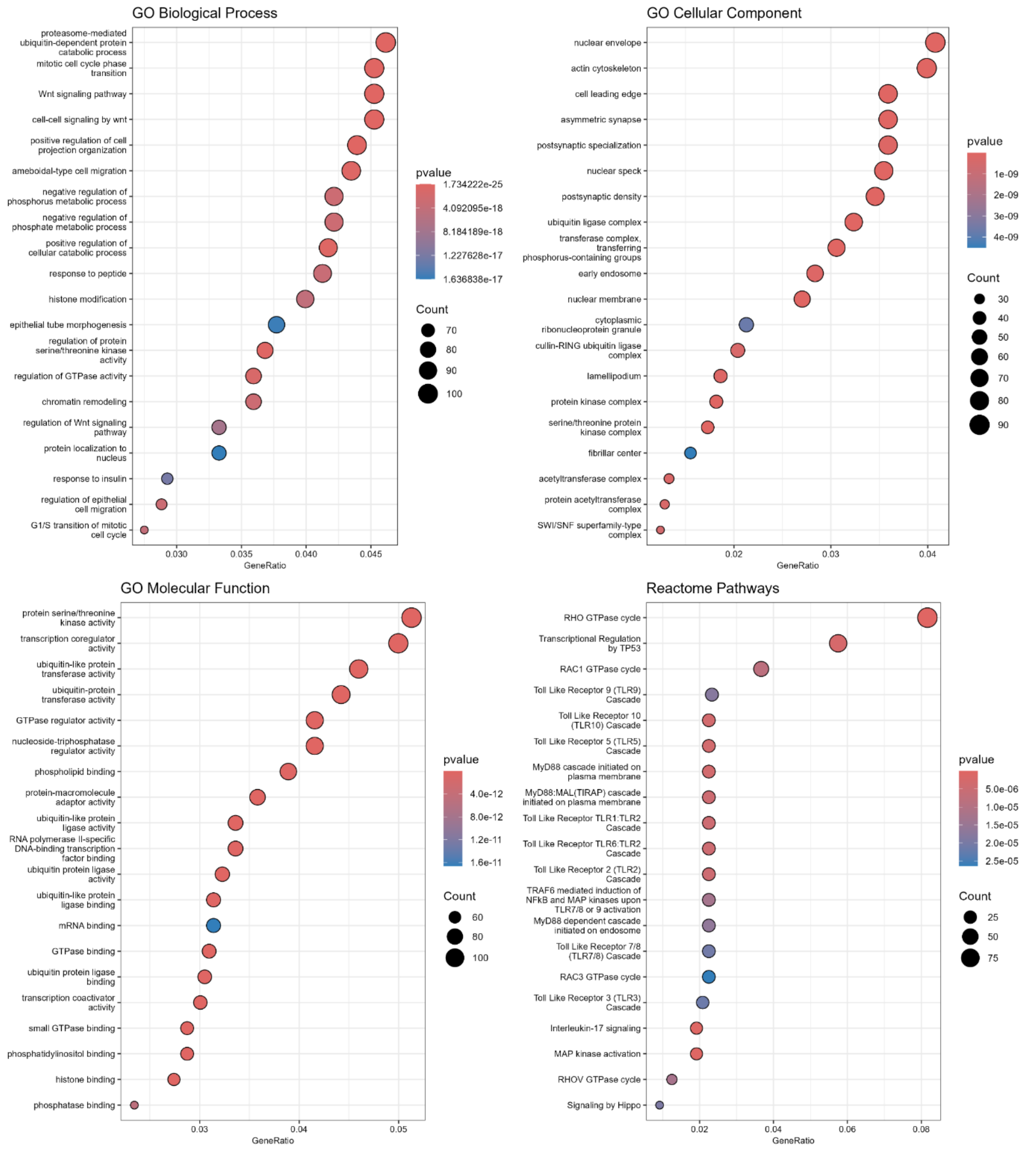


**Figure S4.** Functional overrepresentation analysis for identified mRNA targets of downregulated miRNA in hippocampus identifying biological processes, cellular components, molecular function and reactome pathways regulated by downregulated miRNA ($p < 0.05$).

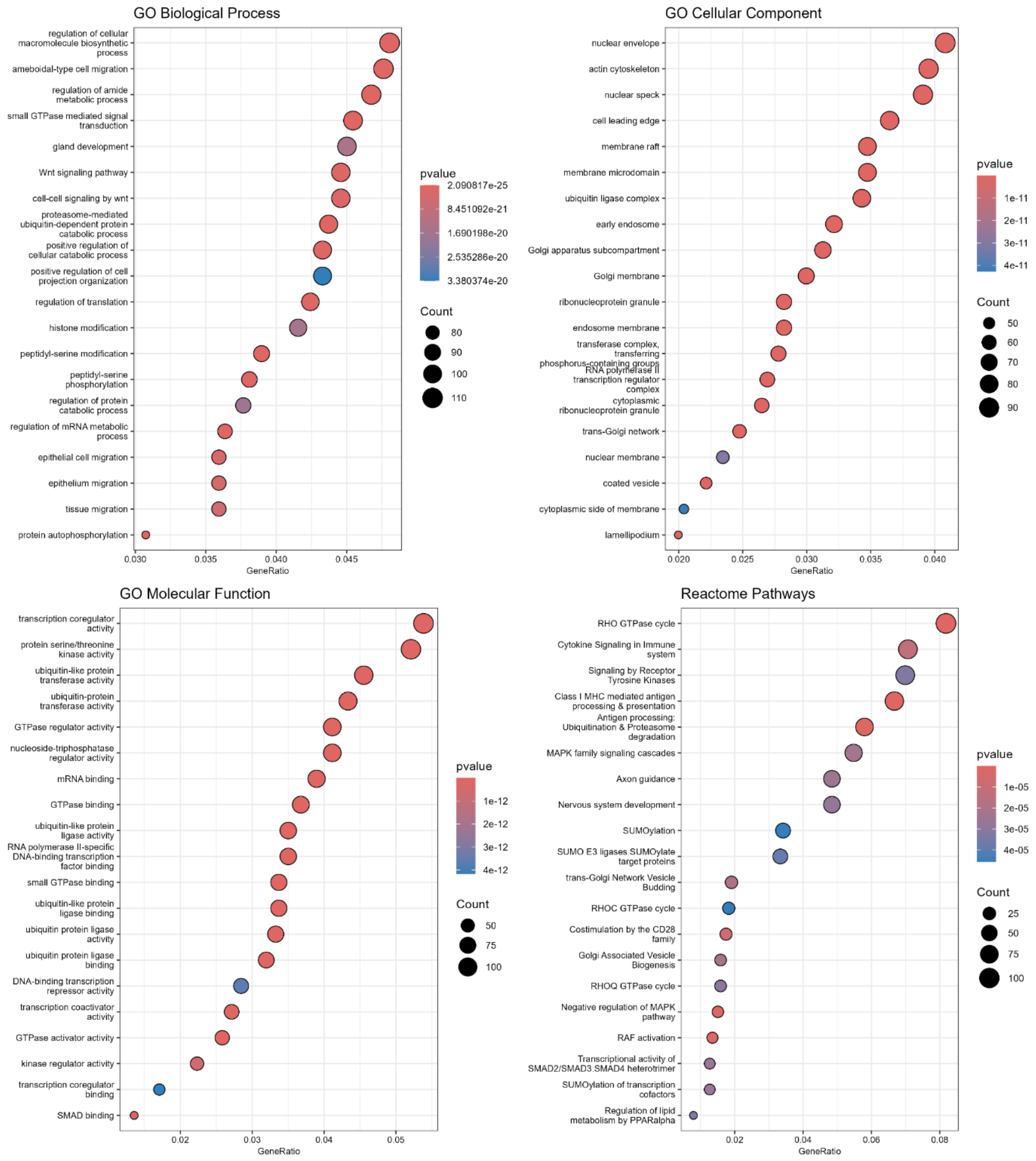


**Figure S5.** Functional overrepresentation analysis for identified mRNA targets of upregulated miRNA in cortex identifying biological processes, cellular components, molecular function and reactome pathways regulated by upregulated miRNA ($p < 0.05$).

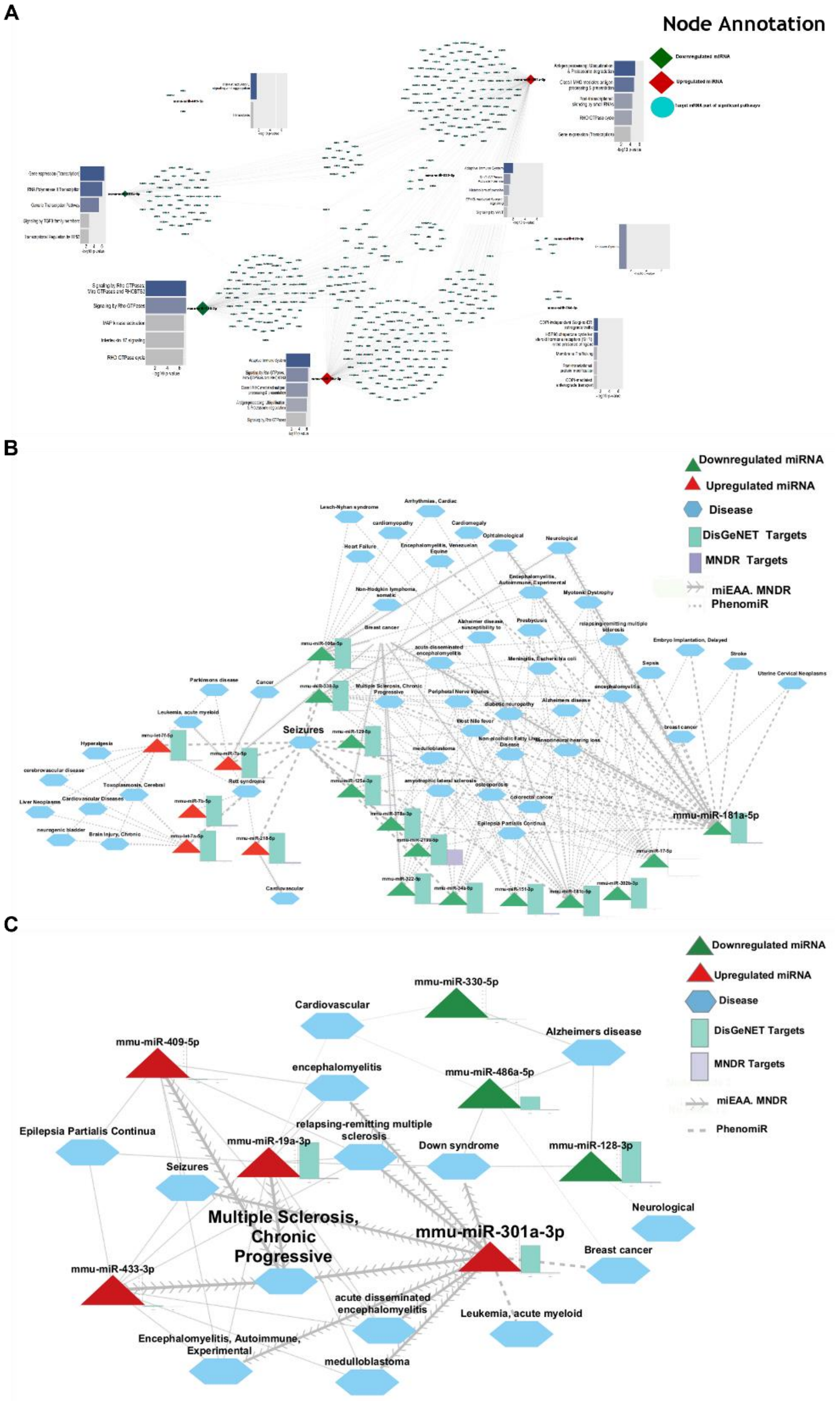


**Figure S6.** (**A**) MiRNA-mRNA-Pathway network constructed for dysregulated miRNA of hippocampus by using top significantly (p value < 0.05) enriched pathways and their corresponding genes. MiRNA-mRNA-disease network constructed for dysregulated miRNA in cortex (**B**) and hippocampus (**C**). Diseases with a p-value <0.05 were considered strongly enriched for the selected collection of miRNAs.

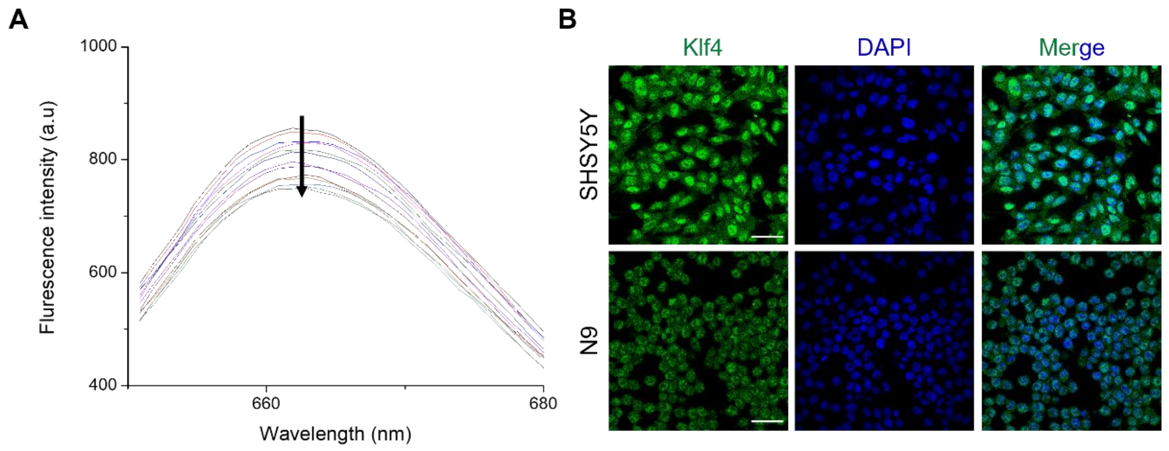


**Figure S7.** (**A**) Fluorescence spectra of Atto-647 labelled miR-7a titrated with Klf4 mRNA complement sequence. (**B**) Representative confocal images Klf4 immunofluorescence in SH-SY5Y and N9 cells.

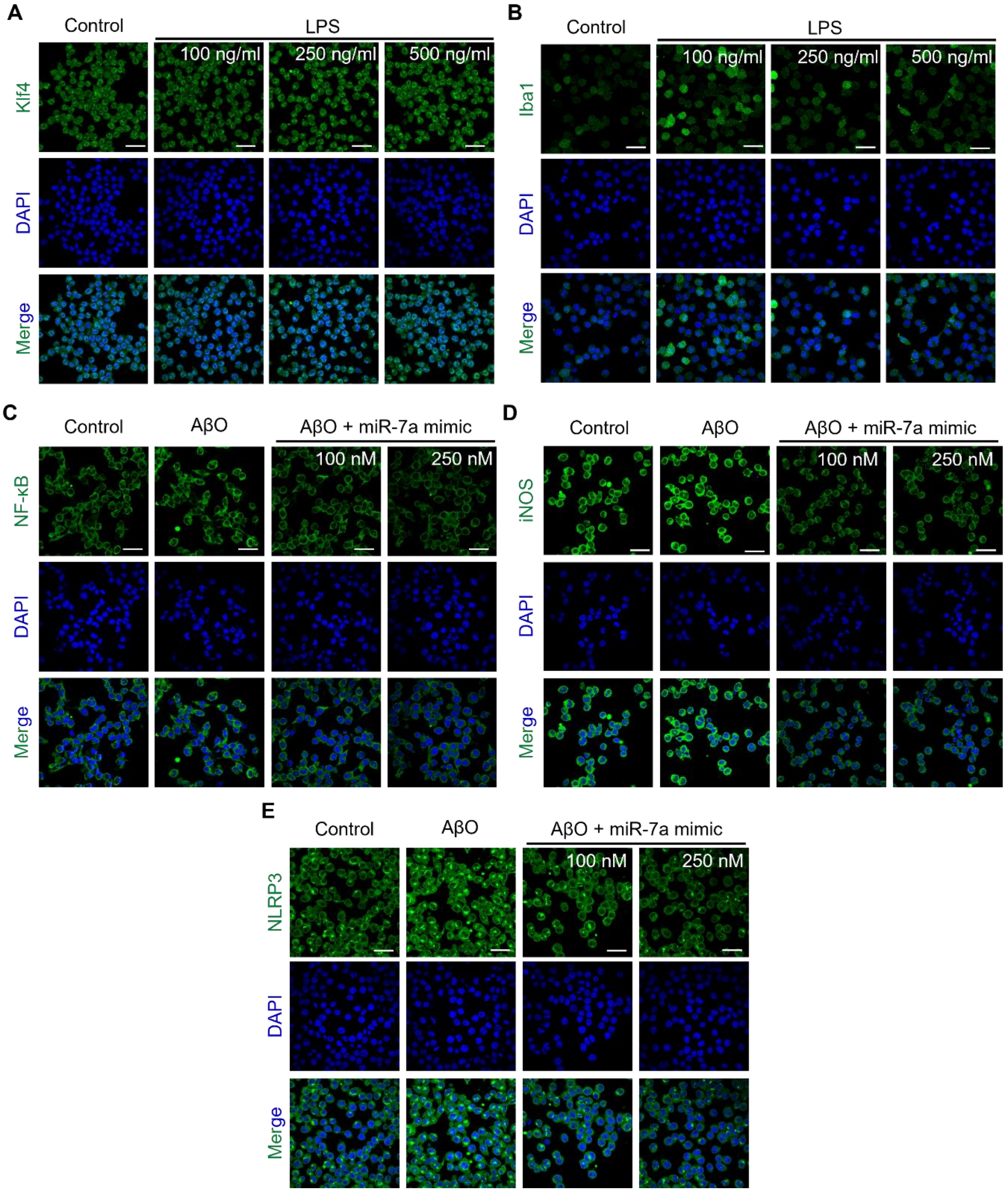


**Figure S8.** Representative confocal images of Klf4 (**A**) and Iba1 (**B**) immunofluorescence in N9 cells treated with increasing concentration of LPS (100, 250 and 500 ng/ml) (Scale bar 200 μM). Representative confocal images NF-κB (**C**), iNOS (**D**) and NLRP3 (**E**)immunofluorescence in N9 cells transfected with miR-7a and challenged with AβO (Scale bar 200 μM).

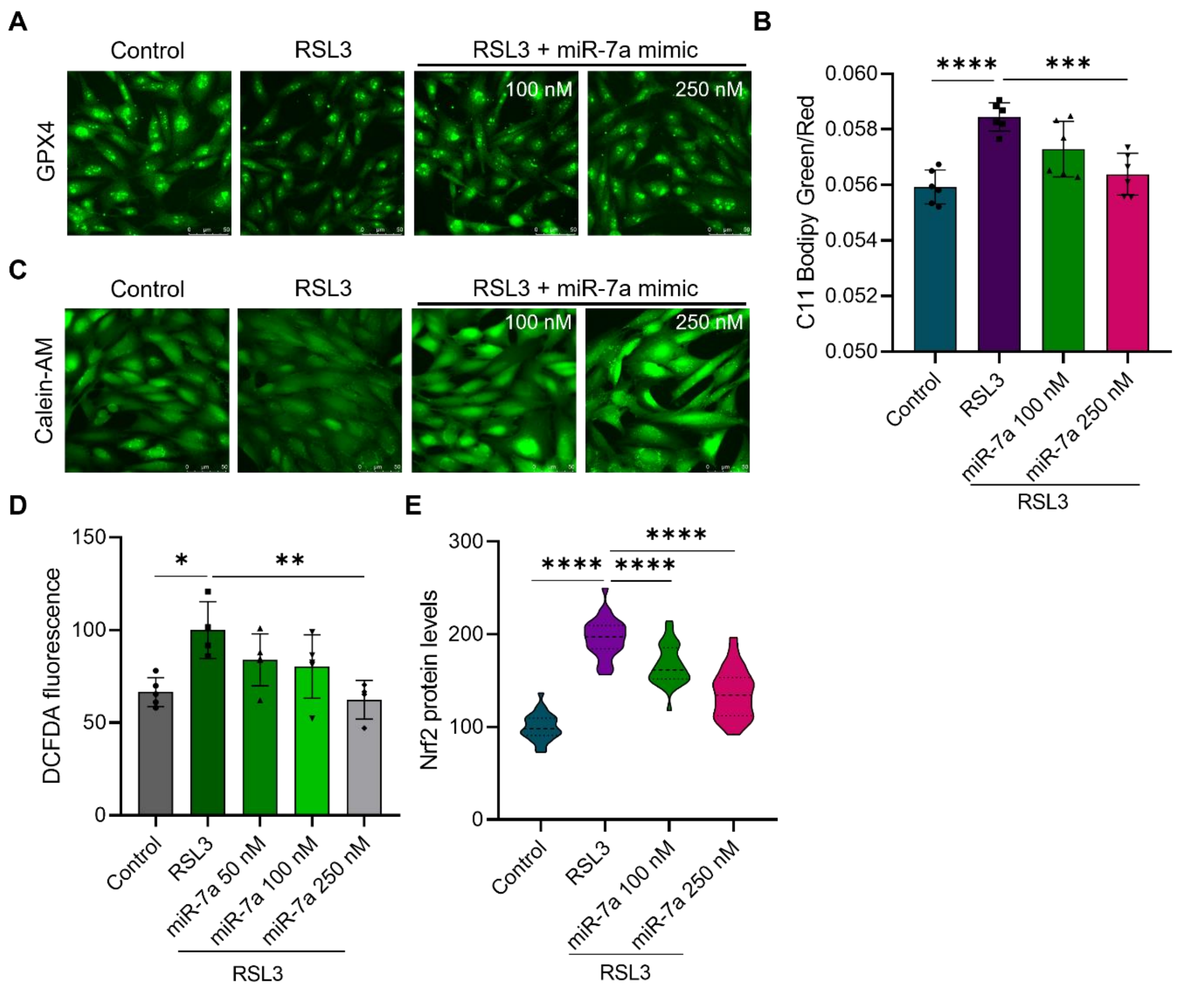


**Figure S9.** (**A**) Representative fluorescence images of GPX4 immunofluorescence in SH-SY5Y cells treated with miR-7a and challenged with RSL3 (Scale bar 50 µM). (**B**) Quantification of lipid peroxidation using C11-Bodipy fluorescent probe in in SH-SY5Y cells treated with miR-7a and challenged with RSL3 in well plate (The bar denotes the ratio of fluorescence intensity at 510 and 591 nm detected by well scan with > 80 data points. n = 6 biological replicates. One-way ANOVA with Bonferroni post hoc test *p < 0.05). (**C**) Representative fluorescence images of calcein-AM in SH-SY5Y cells treated with miR-7a and challenged with RSL3 (Scale bar 50 µM). (**D**) Relative intracellular ROS measured by DCFDA fluorescence (The bar denotes the normalized mean DCFDA fluorescence detected by well scan with > 80 data points. n = 4 biological replicates. One-way ANOVA with Bonferroni post hoc test *p < 0.05). (**E**) Quantification of relative Nrf2 levels in RSL3 induced ferroptotic cells and rescue by miR-7a mimic (n > 500 cells quantified for each group. One-way ANOVA with Bonferroni post hoc test *p < 0.05).

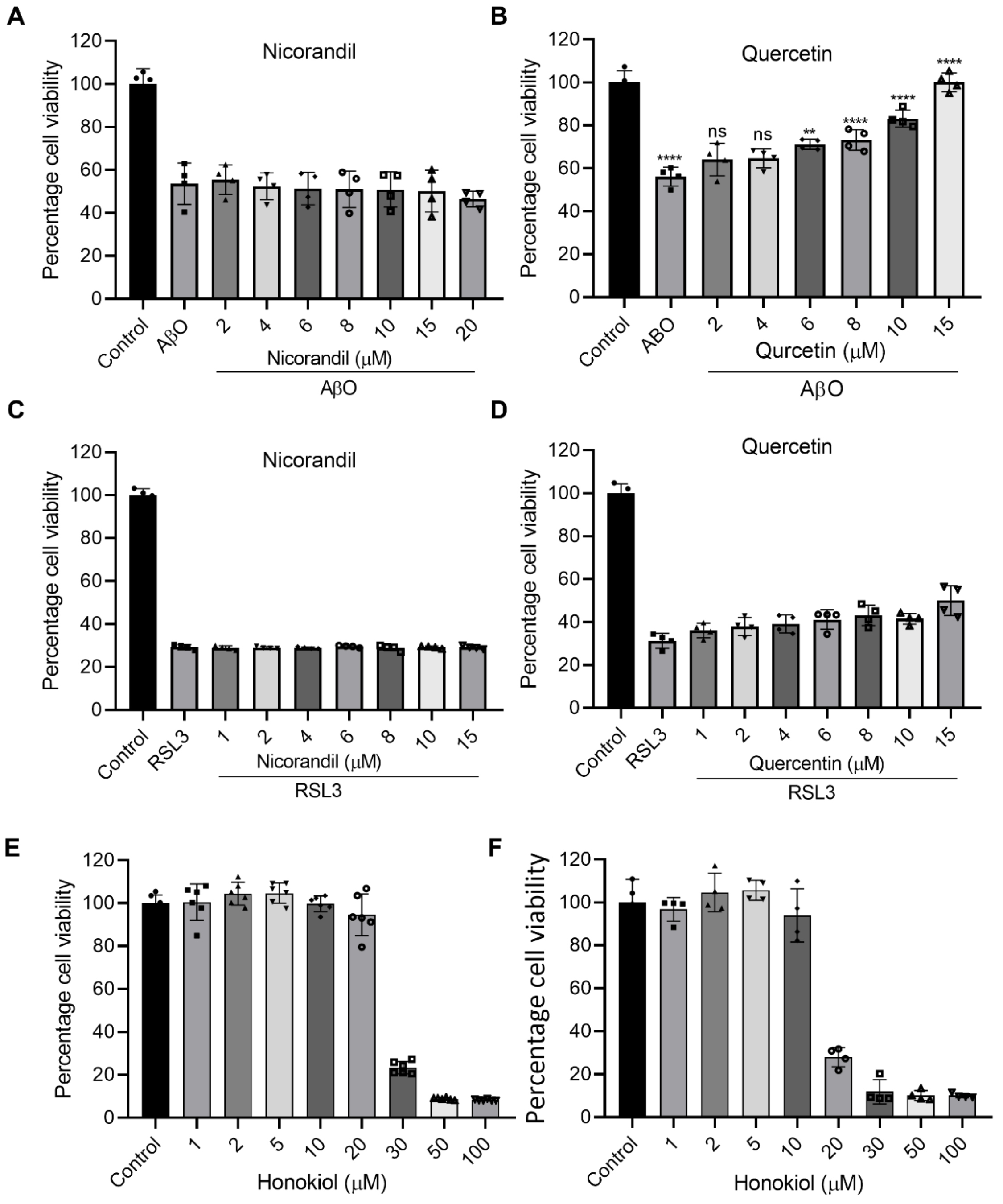


**Figure S10.** N9 microglial cell rescue from AβO induced inflammatory cell death by nicorandil (**A**) and quercetin (**B**) in dose dependent manner (n = 8 biological replicates. One-way ANOVA with Bonferroni post hoc test $^*p < 0.05$). SH-SY5Y neuronal cell rescue from RSL3 induced ferroptotic cell death by nicorandil (**C**) and quercetin (**D**) in dose dependent manner (n = 8 biological replicates. One-way ANOVA with Bonferroni post hoc test $^*p < 0.05$). The cell viability of (**E**) SH-SY5Y and (**F**) N9 cells treated with different concentrations of honokiol (1 to 100 μM) by MTT assay.

**Table S1.** List of antibodies and information used in the study.

| Sl. No | Antibody | Origin | Reactivity | Dilution (IF/ western) | Catalogue | Company |
|---|---|---|---|---|---|---|
| 1 | Klf4 (6C5) mAb | Rabbit | Human, mouse and rat | 1:200/1000 | BSM-52850R | Invitrogen |
| 2 | Nrf2 pAb | Rabbit | Human, mouse and rat | 1:200 | PA5-88084 | Invitrogen |
| 3 | Iba1(E404W) mAb | Rabbit | Human, mouse, rat, hamster, Monkey | 1:200 | 17198S | CST |
| 4 | Iba1 | Chicken | Human, mouse, rat and ape | 1:200 | 234 009 | Synaptic systems |
| 5 | TNFα (D2D4) mAb | Rabbit | Mouse | 1:200 | 11948S | CST |
| 6 | NF-κB (ARC0084) mAb | Rabbit | Human, mouse and rat | 1:200 | A19605 | ABclonal |
| 7 | NLRP3 pAb | Rabbit | Human, mouse and rat | 1:200 | A12694 | ABclonal |
| 8 | iNOS pAb | Rabbit | Human, mouse and rat | 1:50 | PA1-036 | Invitrogen |
| 9 | GPX4 pAb | Rabbit | Human, mouse and rat | 1:200 | E-AB-64550 | Elabscience |
| 10 | Anti-Rabbit IgG (H+L) Highly Cross-Adsorbed Secondary Antibody, Alexa Fluor™ 488 | Goat | Rabbit IgG | 1:500 | A-11034 | Invitrogen |
| 11 | Anti-Chicken IgY (H+L) Highly Cross-Adsorbed Secondary Antibody, Alexa Fluor™ 568 | Goat | Chicken IgG | 1:500 | A-11041 | Invitrogen |

**Data S1. (separate file)** Cortex miRNA-MRNA-pathway network.

**Data S2. (separate file)** Hippocampus miRNA-MRNA-pathway network.

**Data S3. (separate file)** Western blots.

**Table S2. (separate file)** Cortex miRNA Transcriptomic data.

**Table S3. (separate file)** Hippocampus miRNA transcriptomic data.

**Table S4. (separate file)** Cortex validated mRNA targets.

**Table S5. (separate file)** Hippocampus validated mRNA targets.

**Table S6. (separate file)** Disgenet cytoscape disease association.

**Table S7. (separate file)** miEAA database disease association.

**Table S8. (separate file)** MNDR disease association.

**Table S9. (separate file)** Phenomir Disease miRNAs cortex and hippocampus.

**Table S10. (separate file)** List of primers and RNA oligos used the study
